# Unlocking consumer coexistence: The role of self-organised spatial heterogeneity

**DOI:** 10.1101/2023.12.05.570113

**Authors:** Christian Guill, Felix Nößler, Toni Klauschies

## Abstract

In metacommunities, habitat heterogeneity facilitates species coexistence if superior competitors disperse maladaptively towards unfavourable habitats or if they hedge insufficiently against fluctuating environmental conditions. We show that similar mechanisms also operate in metacommunities with homogeneous habitat quality when heterogeneous biomass distributions emerge from self-organised pattern formation. Depending on whether the induced biomass patterns are static or fluctuating, either lower or higher dispersal rates can allow inferior competitors to coexist with their superior counterparts. Coexistence is further promoted when the inferiors can plastically reduce emigration from resource-rich patches. Furthermore, if the competitors differ in their abilities to induce pattern formation, a novel coexistence mechanism akin to relative non-linearity emerges, where the temporarily dominant competitor modifies the spatio-temporal variation in the biomass distributions such that it favours the recovery of the currently rare competitor. Self-organised pattern formation thus generically provides mechanisms for maintaining diversity in metacommunities without requiring a priori habitat heterogeneity.

## 1 Introduction

Understanding the processes that determine species coexistence in natural ecosystems is central to community ecology (Hutchinson 1961; Chesson 2000; Levine et al. 2017). Classical theory distinguishes between mechanisms that operate on the local versus the regional scale of metacommunities, i.e., communities of habitat patches that are linked by dispersal of various species (Leibold et al. 2004). Local coexistence mechanisms include relative nonlinearity in resource-dependent growth functions (e.g. the gleaner-opportunist trade-off, Litchman and Klausmeier 2001; Klauschies and Gaedke 2020), resource partitioning (Schoener 1974), and the temporal storage effect (Chesson and Warner 1981). In contrast, dispersal between different habitats is a key process for coexistence of species in metacommunities that may not be able to coexist in a non-spatial context (Schlägel et al. 2020).

Several mechanisms have been proposed that can explain species coexistence through dispersal in metacommunities. First, the competitive abilities of species may be strongly influenced by abiotic factors or by biotic interactions with other species, leading to a situation where each competitor has at least one patch where it is dominant. The species thus have different source and sink patches (with positive or negative population growth, respectively) and can persist in all habitat patches due to dispersal from source to sink patches (Shmida and Wilson 1985; Amarasekare 2010)). Other coexistence mechanisms assume that the competitive rankings are spatially homogeneous, i.e., source and sink patches are the same for all species. If patch quality is constant in time, coexistence is possible if the dispersal rate of a superior competitor (specifically, its emigration rate from source into sink patches) is higher than that of an inferior competitor (Abrams and Wilson 2004; Namba and Hashimoto 2004; Nathan et al. 2013). Conversely, if the resource availability on the different patches changes over time, coexistence is facilitated if the inferior competitor has a higher dispersal rate than the superior one, similar to the classic competition-colonisation trade-off (Lin et al. 2013).

A prerequisite for coexistence of an inferior competitor with a superior one that is dominant in all patches is that resource densities are not identical everywhere, as otherwise the superior competitor would always competitively exclude the inferior one (Hardin 1960). Usually it is assumed that the patches have different habitat quality due to heterogeneous environmental conditions (Amarasekare 2010; Lin et al. 2013). However, both static and temporally varying patterns in the quality of the patches can also emerge in a self-organised way, and several studies have demonstrated how this promotes species coexistence and biodiversity (Banerjee and Petrovskii 2011; Baudena and Rietkerk 2012; Nathan et al. 2013; Eigentler 2021; Guill et al. 2021). These patterns are formed by an interplay between the local interactions of the species, which often involve some form of system-specific ecosystem engineering process, and their spatial dispersal dynamics (scale-dependent feedback, Rietkerk and Van de Koppel 2008). Because the magnitude and type (static or oscillatory) of the emergent biomass patterns depend on the relative abundance of the coexisting species and their dispersal rates, a feedback loop between the cause and the consequences of the mechanisms underlying coexistence is created, which is likely to affect the range of conditions for their applicability.

Conventionally, studies on self-organised pattern formation in ecological systems have only considered random (diffusive) dispersal. However, coexistence of species in spatially extended systems may also depend on their ability to plastically adjust their dispersal rates depending on, e.g., food availability or predation risk (Bowler and Benton 2005; Schlägel et al. 2020). Indeed, it has often been shown that plastic responses or rapid trait evolution, as it appears in e.g. adaptive foraging (Heckmann et al. 2012) or defence against predators (van Velzen 2020), can enable coexistence where it is not possible with fixed traits. Corresponding plasticity of dispersal strategies not only implies that dispersing individuals migrate towards patches with favourable conditions, but also that their emigration from these patches is suppressed (Amarasekare 2010; Cressman and Křivan 2013; Lin et al. 2013).

In this study we combine these approaches to address how different dispersal rates and levels of dispersal plasticity affect species coexistence through their influence on self-organised formation of heterogeneous biomass patterns. In order to obtain widely applicable results, the considered system is deliberately kept very simple. On two adjacent patches with identical environmental conditions, two competing species (heterotrophs) with different competitive strength feed on a shared resource (autotrophs), which in turn relies on nutrients (Fig. 1a-c). In addition to movement of all constituents between the patches, the model thus only contains generic trophic and (indirect) competitive interactions and does not invoke any ecosystem engineering processes to induce self-organised pattern formation. In such a setting, the dispersal rates of the competitors are important determinants of whether no, static, or oscillatory patterns in the species’ biomass distributions emerge (Guill et al. 2021). At the same time, the relative dispersal rates of the competitors contribute directly to their fitness in heterogeneous metacommunities. We show that this, in combination with the feedback between pattern formation and species abundances, leads to coexistence under conditions where it would not be possible with externally determined environmental heterogeneity.

**Figure 1.**
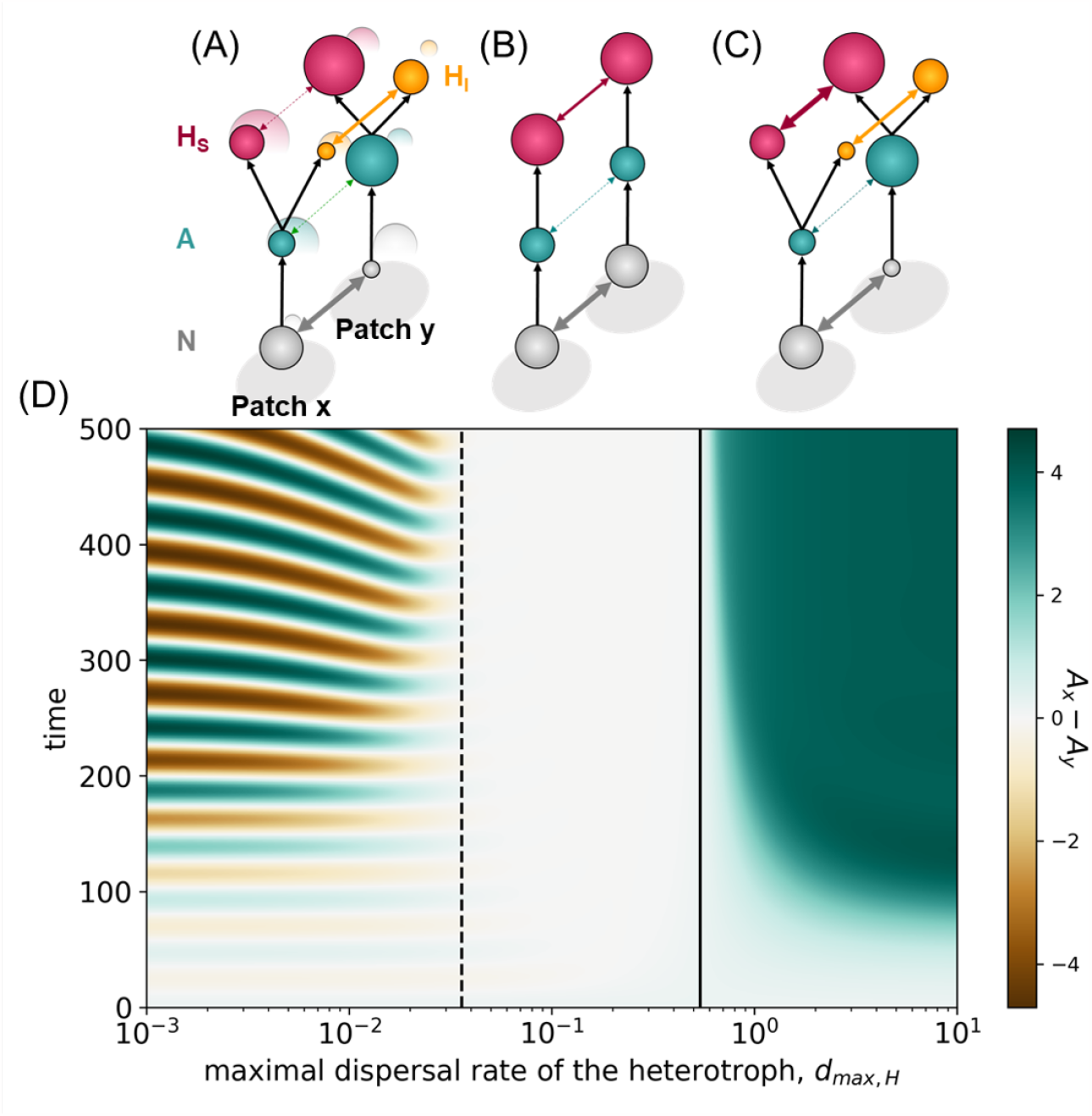
Conceptual representation of the model. The system contains nutrients (*N*, grey), autotrophs (*A*, green), and two competing heterotrophs (superior competitor *H*_*S*_ in red, inferior competitor *H*_*I*_ in orange) in the two habitat patches *x* and *y*. Black arrows denote local trophic interactions, coloured arrows indicate dispersal. The size of the spheres represents the nutrient and population densities in the respective habitat patch. The dispersal rate of the superior competitor determines the basic form of biomass patterns emerging: A) oscillatory pattern formation at low dispersal rate, B) no pattern formation (and exclusion of the inferior competitor) at intermediate dispersal rate, and C) static pattern formation at high dispersal rate. In panel D), time series of the difference in autotroph density between the two patches for different heterotroph dispersal rates are shown (with only one heterotroph in the system). The dashed vertical line indicates an oscillatory Turing instability, the solid vertical line indicates a static Turing instability. See Table 1 for parameter values.

## 2 Methods

To analyse the conditions under which two competitors with different dispersal and competitive abilities can coexist in a system of two identical habitat patches we developed a metacommunity model that describes the local dynamics of a simple food web comprising nutrients (with concentration *N*), one autotroph and two heterotroph species (with biomass densities *A, H*_*S*_, and *H*_*I*_, respectively) in each patch and the diffusion of nutrients and dispersal of individuals between the two patches.

**Table 1:** Description, dimensions and values of the model parameters. The parameters without a value are varied and in that case, the values are explicitly mentioned in the text and below the figures. The subscript *i* ∈ {*S, I* } refers to either the superior or the inferior competitor.

| Name | Description | Dimension | Value |
| --- | --- | --- | --- |
| $S$ | nutrient supply concentration | $\text{mass} \cdot \text{area}^{-1}$ | 4.8 |
| $D$ | dilution rate | $\text{time}^{-1}$ | 0.3 |
| $N_h$ | half-saturation constant for nutrient uptake | $\text{mass} \cdot \text{area}^{-1}$ | 1.5 |
| $r_{max}$ | maximal uptake rate of autotroph | $\text{time}^{-1}$ | 0.7 |
| $a_i$ | attack rates of the heterotrophs | $\text{area} \cdot (\text{mass} \cdot \text{time})^{-1}$ | 1.2, 1.0 |
| $h$ | handling time of the heterotrophs | time | 0.53 |
| $e$ | conversion efficiency of the heterotrophs | dimensionless | 0.33 |
| $d_N$ | dispersal rate of nutrients | $\text{time}^{-1}$ | 1.0 |
| $d_A$ | dispersal rate of autotroph | $\text{time}^{-1}$ | 0.001 |
| $k_i$ | sensitivity of the heterotrophs | $\text{area} \cdot \text{mass}^{-1}$ | - |
| $d_{max,i}$ | maximal dispersal rate of the heterotrophs | $\text{time}^{-1}$ | - |

### 2.1. Model description

The differential equations describing the population dynamics of the food web on the first patch (with index *x*) are as follows:

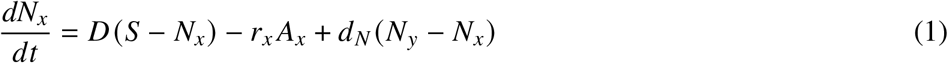

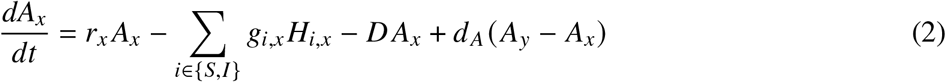

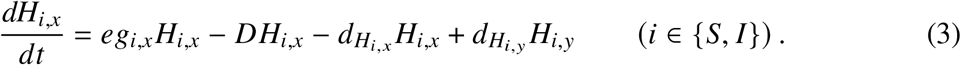

The equations for the second patch (with index *y*) are obtained symmetrically by swapping the indices *x* and *y*. All model parameters are summarised in Table 1, a flow diagram of the model is provided in Fig. S1.

Locally, the model describes flow-through systems like chemostats. Nutrient-rich medium with supply concentration *S* is constantly replenished with turnover rate *D*, which also determines the per capita mortality rate of autotrophs and heterotrophs. The nutrient uptake rate of the autotroph, *r*_*x*_, follows Michealis-Menten kinetics (Michaelis and Menten 1913),

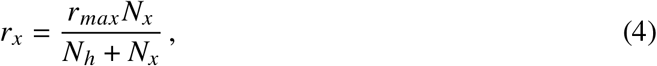

with maximum uptake rate *r*_*max*_ and half saturation concentration *N*_*h*_. The grazing rates of the heterotrophs, *g*_*i,x*_ are modelled as Holling type II functional responses (Holling 1959),

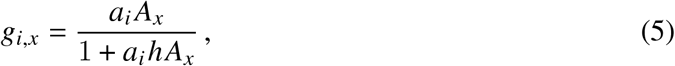

with attack rates *a*_*i*_ (which determine the competitive hierarchy of the heterotrophs, i.e., *a*_*S*_ *> a*_*I*_) and handling time *h*. Consumed autotroph biomass is converted to heterotroph biomass with conversion efficiency *e*, accounting for fecal and respiratory energy loss.

Nutrients and autotrophs are assumed to randomly diffuse or disperse between the patches with constant per capita rates *d*_*N*_ and *d*_*A*_, respectively. In contrast, it is assumed that the per capita dispersal rates of the heterotrophs can plastically respond to the prevailing local food conditions following a reversed logistic function:

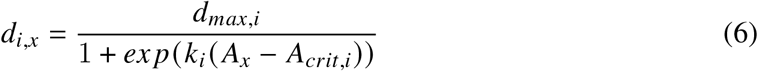

with maximal dispersal rate *d*_*max,i*_ and sensitivity *k*_*i*_. The inflection point *A*_*crit,i*_ is given by the autotroph density at which the local per capita growth rate of the respective heterotroph is zero:

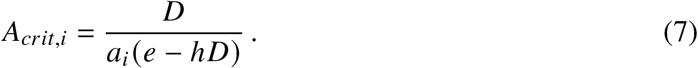

If the local autotroph density is higher than *A*_*crit,i*_, the growth rate of heterotroph *i* is positive and its dispersal rate out of patch *x, d*_*i,x*_, is less than half the maximal dispersal rate, otherwise it is higher. The sensitivity *k*_*i*_ determines the steepness of the transition around *A*_*crit,i*_. If *k*_*i*_ = 0, the heterotroph disperses randomly with a constant per capita dispersal rate of half its maximal dispersal rate. Note that we do not assume a colonisation-competition trade-off in the parametrisation of the model, i.e., it is not required that the competitively inferior heterotroph *H*_*I*_ has a higher maximum dispersal rate or sensitivity than the competitively superior heterotroph *H*_*S*_.

### 2.2. Model analysis

First we determined the conditions for self-organised pattern formation. Since the onset of this phenomenon is caused by a Turing instability of a stable, spatially homogeneous state (Turing 1952), which does not allow for coexistence of the two heterotrophs (Appendix S2), the analyses were carried out in a system with just a single heterotroph. As important parameters affecting the pattern formation process, the dispersal rates *d*_*N*_, *d*_*A*_, and *d*_*max,H*_, the sensitivity *k*_*H*_, and the attack rate *a*_*H*_ were included in the analyses (note that we use the subscript *H* instead of *i* to distinguish the model setup with just a single heterotroph from the standard case with two competing species). The procedures for determining the location of the Turing instability in parameter space are detailed in Appendix S3. Provided that self-organised pattern formation can occur, further coexistence conditions were then explored in the full system with both competitors. For this, the effect of the maximal dispersal rates *d*_*max,i*_ was studied in detail for two sensitivity levels of the inferior competitor (*k* _*I*_ = 0 and *k* _*I*_ = 2). Additional analyses on the effect of the attack rates *a*_*i*_ and sensitivities *k*_*i*_ are presented in Appendix S4.

In order to test for coexistence of the two competitors, the system was simulated first with only the superior competitor, *H*_*S*_, for 10^4^ time steps. Then, the inferior competitor, *H*_*I*_, was added with a low density (*H*_*I,x*_ = 0.001, *H*_*I,y*_ = 0.0001). The system was simulated for another 10^5^ time steps. If at the end of a simulation run both competitors had a density greater than 10^−10^ summed over both patches, coexistence was assumed. For numerical reasons, a variable was set to 0 if its value dropped below 10^−30^ during a simulation run. We checked that our results are largely independent of the order of invasions (Appendix S4).

The simulations were performed in Julia 1.9.3 (Bezanson et al. 2017) with the help of the packages DifferentialEquations.jl (Rackauckas and Nie 2017). The numerical solver Vern9 (Verner 1978) with an absolute and relative tolerance of 10^−12^ from the DifferentialEquations.jl package was used. To check that the choice of the solver had no substantial effect on the results, some simulations were repeated with the CVODE solver (Hindmarsh et al. 2005). The figures were produced with Makie.jl (Danisch and Krumbiegel 2021). We verified the reproducibility of the results using Python simulations. Most of the supporting figures were produced with Python using the packages SciPy, NumPy and Matplotlib. The simulation code and detailed information on the required versions of the Julia and Python packages used is available online (see data accessibility statement).

## 3 Results

### 3.1. Self-organised pattern formation

In the absence of self-organised pattern formation, the superior competitor always outcompetes the inferior one as the patches have the same a-priori habitat quality and the inferior competitor cannot invade into a homogeneous system with resident superior competitor, while the reverse is always possible (Appendix S2).

The onset of pattern formation is marked by a spatial instability (Turing instability) of a homogeneous single-heterotroph equilibrium. Depending on the parameters of the system (most importantly the maximum dispersal rate of the heterotroph, *d*_*max,H*_) either static or oscillatory (spatio-temporal) patterns can occur. At low values of *d*_*max,H*_, an oscillatory Turing instability occurs (marking the emergence of spatio-temporal patterns), while at high values of *d*_*max,H*_, static patterns emerge (Fig. 1D). At intermediate levels of *d*_*max,H*_, the homogeneous equilibrium remains stable. Under plastic dispersal of the heterotroph, only low values of the sensitivity (*k*_*H*_ ≲ 0.4) allow for the emergence of static patterns at high values of *d*_*max,H*_ (Fig. S4). In contrast, the emergence of oscillatory patterns is not affected by dispersal plasticity. Further requirements for pattern formation are that the diffusion rate of the nutrients is sufficiently high (*d*_*N*_ ≳ 0.1) and the dispersal rate of the autotrophs is comparatively small (*d*_*A*_ ≲ 0.03, Fig. S4).

### 3.2. Coexistence patterns and mechanisms

In Fig. 2, coexistence of a superior competitor, *H*_*S*_, with an inferior one, *H*_*I*_, is shown as a function of their maximal dispersal rates *d*_*max,S*_ and *d*_*max,I*_. For intermediate values of *d*_*max,S*_ (grey area) no Turing instability is induced and coexistence is therefore not possible. At low values of *d*_*max,S*_ (≲ 0.05) an oscillatory Turing instability is induced and the autotroph densities on the two patches cycle in anti-phase (see also Fig. 3A). Coexistence is possible if *H*_*I*_ has a (moderately) higher dispersal rate than *H*_*S*_. While the increase of *H*_*S*_ on patch *x* (between the red triangle markers in Fig. 3A) is almost completely driven by autochthonous growth, *H*_*I*_ starts to accumulate biomass on this patch even before the autotroph density is high enough for a positive growth rate (orange upright triangle marker in Fig. 3A). Essentially, the higher mobility enables *H*_*I*_ to shift biomass from a patch with momentarily favourable, yet declining growth conditions into a patch with momentarily unfavourable, yet improving growth conditions, suggesting that coexistence is based on bet-hedging behaviour of *H*_*I*_. Coexistence is not possible if *d*_*max,I*_ is very high, as this moves too much biomass of *H*_*I*_ from the (temporary) source patch into the (temporary) sink patch (Fig. 2A). Only if *H*_*I*_ can plastically reduce its dispersal rate when growth conditions are good coexistence is possible with arbitrarily high *d*_*max,I*_ (Fig. 2B).

**Figure 2.**
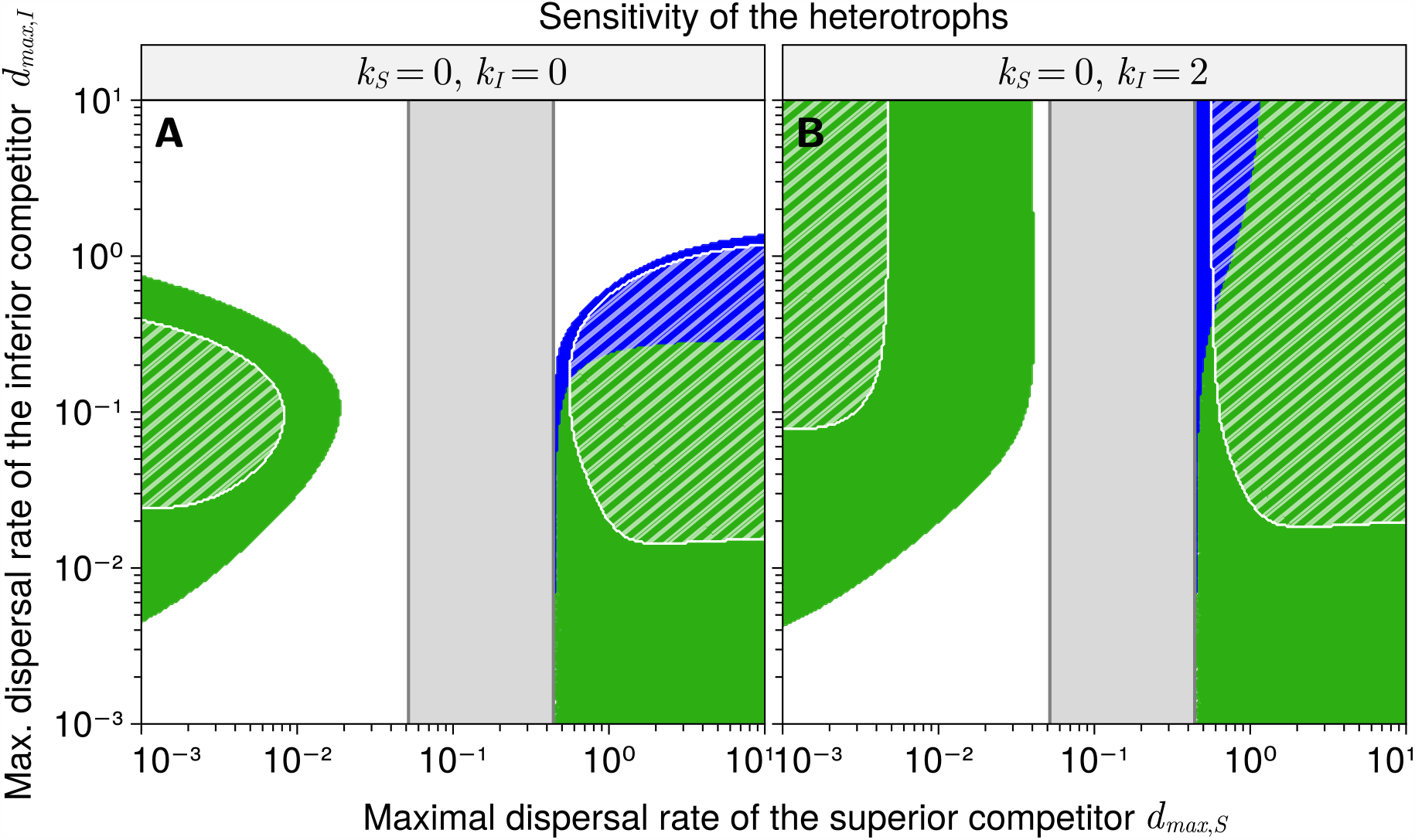
Coexistence as a function of the maximal dispersal rates of the competing heterotrophic species (A: random dispersal of both competitors, B: plastic dispersal of the inferior competitor). Self-organised pattern formation allows for coexistence in the coloured regions (blue: static coexistence, green: oscillatory or chaotic dynamics). In the grey area the superior competitor *H*_*S*_ does not induce a Turing instability. In the hatched areas the inferior competitor *H*_*I*_ excludes *H*_*S*_ if habitat heterogeneity is determined by the environment instead of emerging in a self-organised way.

**Figure 3.**
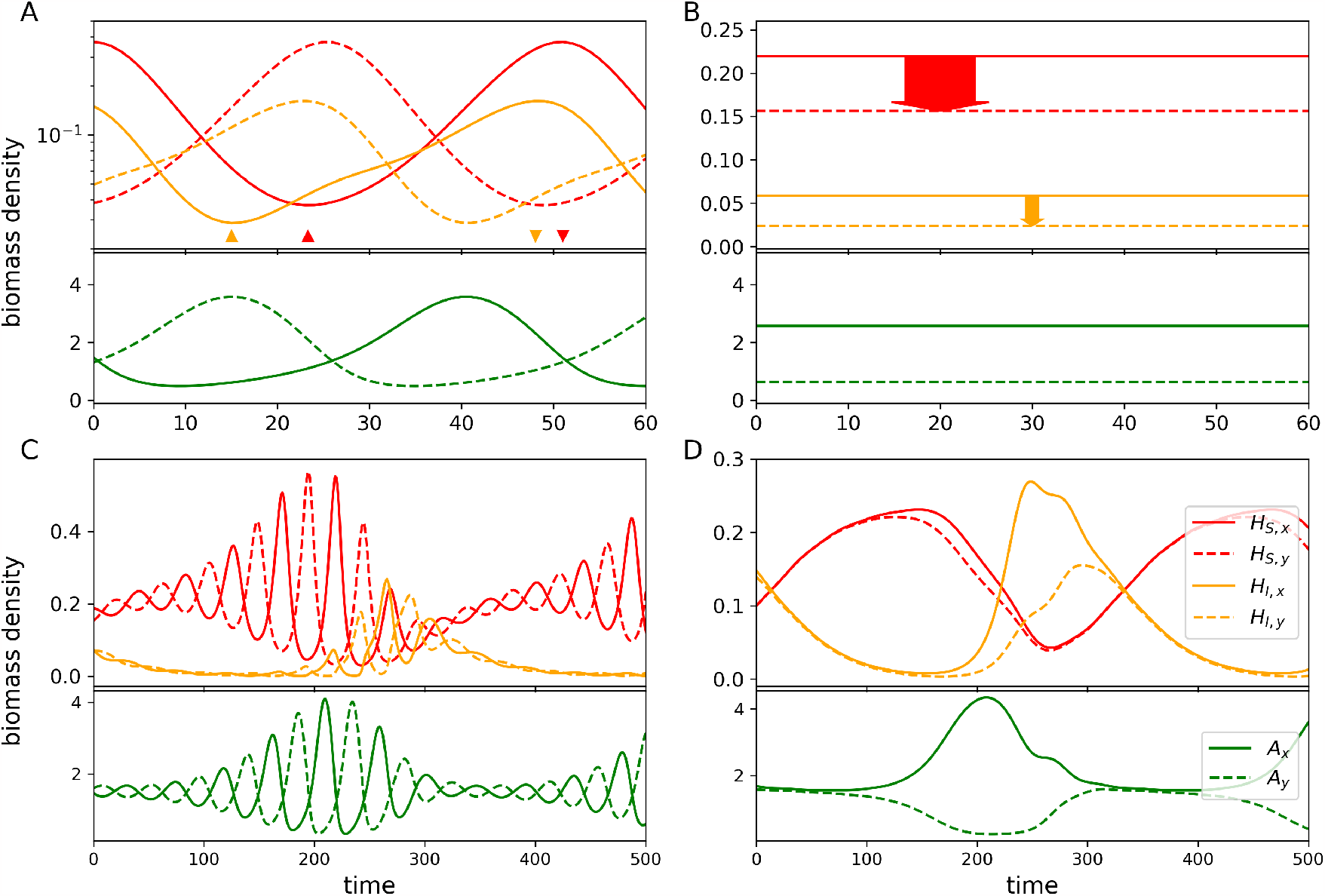
Time series of autotroph (green) and heterotroph densities (*H*_*S*_ in red, *H*_*I*_ in orange) illustrating the coexistence mechanisms. Solid lines: densities on patch *x*, dashed lines: densities on patch *y*. A: bet-hedging mechanism, *k* _*I*_ = 0, *d*_*max,S*_ = 0.01, *d*_*max,I*_ = 0.1. The markers indicate where the the competitors start (stop) accumulating biomass on patch *x*. B: maladaptive-dispersal mechanism, *k* _*I*_ = 0, *d*_*max,S*_ = 0.6, *d*_*max,I*_ = 0.2. The width of the arrows is proportional to the net flow of biomass from patch *x* into patch *y*. C: bet-hedging with heterogeneity modulation, *k* _*I*_ = 2, *d*_*max,S*_ = 0.01, *d*_*max,I*_ = 0.2. D: maladaptive dispersal with heterogeneity modulation, *k* _*I*_ = 0, *d*_*max,S*_ = 2, *d*_*max,I*_ = 0.2. Note the different scaling of the axes among panels. In all panels *k*_*S*_ = 0, all other parameters are as in Table 1.

At high values of the maximal dispersal rate of *H*_*S*_ (*d*_*max,S*_ ≳ 0.44), a static Turing instability is induced, characterised by constantly large differences in autotroph density *A* between the patches (Fig. 3B). Under these conditions, dispersal of *H*_*S*_ is very maladaptive, as it means exporting lots of biomass from the source (high *A*) to the sink (low *A*) patch (red arrow in Fig. 3B). Coexistence is possible if *H*_*I*_ has a lower constant dispersal rate than *H*_*S*_ (*k* _*I*_ = 0, Fig. 2A) or if it can plastically reduce the emigration rate from the source patch (*k* _*I*_ = 2, Fig. 2B), either of which balances the competitive disadvantage of *H*_*I*_ by limiting its loss of biomass due to maladaptive dispersal into the sink patch (orange arrow in Fig. 3B). Provided a sufficiently low dispersal rate, *H*_*I*_ can even persist if its attack rate is lower than what is required in homogeneous systems (Fig. S2 and Fig. S7C), pointing at emergent facilitation by its competitor *H*_*S*_.

If pattern formation leads to coexistence of the two heterotroph species, we often observe population oscillations, irrespective of whether *H*_*S*_ initially created a static or an oscillatory spatial instability (Fig. 2, green areas, and Fig. 3C, D). This is because if *H*_*I*_ persists with a significant density, it also affects the process of pattern formation. For many combinations of the maximum dispersal rates *d*_*max,S*_ and *d*_*max,I*_, the advantage of *H*_*I*_ due to its superior dispersal strategy (given the amount of resource heterogeneity created by *H*_*S*_) outweighs its competitive inferiority. As *H*_*I*_ becomes more abundant and starts to replace *H*_*S*_, it suppresses pattern formation or modifies it (from static to oscillatory patterns or vice versa), which causes *H*_*I*_ to lose its advantage again. The time series in Fig. 3 demonstrate the ensuing periodic modulation of heterogeneity in the autotroph densities for both coexistence mechanisms discussed above. Temporary dominance of *H*_*S*_ causes the formation of pronounced spatial patterns (high-amplitude anti-phase oscillations, Fig. 3C, or a large static difference between *A*_*x*_ and *A*_*y*_, Fig. 3D). Under these conditions, a significantly higher or lower dispersal rate, respectively, allows *H*_*I*_ to outcompete *H*_*S*_, which, however, then suppresses the spatial heterogeneity in autotroph density and allows *H*_*S*_ to grow again.

In order to assess the importance of this dynamic modulation of spatial heterogeneity for coexistence, we contrasted the predictions of our model with one that includes the same amount of resource heterogeneity between the patches as *H*_*S*_ creates at a given level of *d*_*max,S*_, but as an environmental factor that is not affected by the population dynamics of the species (see Appendix S5 for details). We found that under these conditions coexistence of consumers with different dispersal strategies is restricted to much narrower parameter ranges, as *H*_*I*_ cannot modify the level of resource heterogeneity and consequently often excludes *H*_*S*_ due its superior dispersal strategy (white-hatched areas in Fig. 2, see also Fig. S8).

Finally, we note that while dispersal plasticity of *H*_*I*_ increases the range of conditions allowing for coexistence if its maximal dispersal rate is high, plastic dispersal of *H*_*S*_ has the opposite effect: it prevents self-organised pattern formation at high maximal dispersal rates (Fig. S4), which makes coexistence impossible (Fig. S6). For both competitors, dispersal plasticity cannot strongly affect the absolute value of their dispersal rate when its maximum is low and therefore also does not affect coexistence.

## 4 Discussion

Using a generic model we show that emergent habitat heterogeneity due to self-organised pattern formation enables coexistence of species with different competitive and dispersal abilities in metacommunities. Coexistence relies either on maladaptive dispersal between emergent source and sink habitats or on bet-hedging in metacommunities with temporally varying resource heterogeneity. Moreover, differences in the species’ potential to induce and benefit from spatial pattern formation create a novel mechanism that allows competing species to coexist under conditions where an equivalent amount of environmental, i.e., exogenously determined, habitat heterogeneity would not.

### 4.1. Consumer coexistence through spatial pattern formation

While superior competitors for shared limiting resources eventually exclude inferior competitors in static, homogeneous environments (Hardin 1960), species coexistence may be possible in the presence of habitat heterogeneity when different dispersal strategies offset differences in the species’ competitive abilities (Abrams and Wilson 2004; Amarasekare 2010). In contrast to most previous studies, we did, however, not assume that habitat heterogeneity is externally determined, but may intrinsically emerge through the interplay of local trophic interactions and dispersal.

Pattern formation in ecological systems has been studied before, most notably in the context of dryland ecosystems (Klausmeier 1999; Kéfi et al. 2010; Meron 2015), but also in other systems such as mussel beds (van de Koppel et al. 2008) or submarine sea grass (Ruiz-Reynés et al. 2017). In these examples, pattern formation is usually linked to an ecosystem engineering process unique to the system under consideration (like the redistribution of ground water by a change in surface water infiltration rate by local vegetation), which limits the transferability of findings to other systems. In contrast, we study a generic two-patch model which, at its core, only comprises a consumer, a resource and a limiting nutrient. This makes it the simplest ecological model in which both oscillatory and static Turing instabilities (the bifurcations leading to self-organised pattern formation) can occur. While most ecological studies on pattern formation so far have focused on primary producers (Borgogno et al. 2009) and its effect on their (functional) diversity (Nathan et al. 2013; Guill et al. 2021), we show here that the phenomenon is also relevant for higher trophic levels.

The basic mechanisms that enable consumer coexistence in this study have been investigated before under the premise of externally provided habitat heterogeneity. In general, coexistence is possible if a trade-off exists between the competitive strength (here expressed as the attack rate on the shared resource) and the dispersal strategy (which includes both the maximal dispersal rate and the ability to plastically adjust the dispersal rate in response to cues like resource availability). What constitutes a superior dispersal strategy, however, is context-dependent. In case of static differences in patch quality, one patch is necessarily a source (enabling positive population growth) while the other is a sink (in which the surplus production of the source dies of) for both species. In this case, a low dispersal rate reduces maladaptive dispersal from source to sink patch and enables coexistence by allowing an otherwise inferior competitor to retain a higher fractions of its total biomass in the source patch (Abrams and Wilson 2004; Namba and Hashimoto 2004; Amarasekare 2010). As is shown here for the coexistence of consumers and by Nathan et al. (2013) for coexistence of terrestrial plants that are limited by ground water, the same mechanism also works when static habitat heterogeneity emerges by self-organisation.

In the contrasting case where the relative quality of the patches changes periodically, an intermediate dispersal rate creates an advantage, which can be best explained through bet-hedging in the spatially and temporally variable landscape. Analogously to bet-hedging strategies that spread reproduction in time via dormant seeds or increased adult survival in temporally variable environments (Childs et al. 2010; Nevoux et al. 2010), dispersal spreads individuals and thus reproductive capacity in space. At intermediate dispersal rates, this increases total population growth rate during times when the population recovers from low density. However, if the maximal dispersal rate is too high and cannot be reduced by plastic behaviour, the phase of positive net immigration into a patch occurs earlier in the population cycle. This implies that a higher fraction of the immigrating individuals faces insufficient resource densities and therefore cannot contribute to population recovery, which ultimately negates the potential of this strategy to balance a competitive disadvantage. Similar coexistence patterns have been found in a predator-prey model with a priory differences in habitat quality (Lin et al. 2013).

Coexistence through bet-hedging depends on temporal fluctuations in the species densities and can thus be viewed as a generalisation of coexistence due to a competition-colonisation trade-off. The latter focuses on the occupancy of patches by competing species (Tilman 1994; Levins and Culver 1971) and considers extreme environmental fluctuations leading to complete extinctions of local populations and necessitating recolonisation of patches. The bet-hedging mechanism in contrast explicitly includes the population dynamics of the competitors and their shared resources in the patches. However, in both cases stability of the environment favours the superior competitor over the species that is dispersing or colonising faster (Pellissier 2015).

An inferior dispersal strategy of the superior competitor may not only enable persistence of the inferior competitor, but can even lead to the exclusion of the superior competitor. This outcome is especially prevalent if the heterogeneous resource distribution results from environmental differences between the patches (e.g., different supply concentration of nutrients). However, if habitat heterogeneity emerges via self-organisation when the superior competitor is dominant, its exclusion due to a disadvantageous dispersal strategy is much less common. As the inferior competitor gains dominance, it modifies the emergent heterogeneous distribution of the resources in a way that makes its dispersal strategy less advantageous (either by dampening the heterogeneity, or by turning static patterns into spatio-temporal patterns or vice versa) and thus allows the superior competitor to recover. This dynamical pattern strikingly resembles coexistence due to a gleaner-opportunist trade-off, where a species that benefits from a fluctuating environment but stabilises it (the opportunist) coexists with a species that benefits from a stable environment but destabilises it (the gleaner, Yamamichi et al. 2022). This mechanism has been considered in a spatial context before (Pacala and Rees 1998; Wilson and Abrams 2005), but only under the premise of emergent or forced temporal fluctuations. In contrast, here we show for the first time that this well-established coexistence mechanism can also build upon dynamical creation and dampening of spatial heterogeneity.

Finally, species coexistence also depends on whether the heterotrophs can plastically adjust their dispersal rates in response to fitness-relevant cues. Previous studies found that such dispersal plasticity is essential for coexistence in fluctuating environments, but may make it impossible in static environments if it allows superior competitors to avoid maladaptive dispersal (Amarasekare 2010; Lin et al. 2013). This contrasts with our results showing that dispersal plasticity is not strictly necessary for coexistence in fluctuating environments and that the ranges of conditions allowing for coexistence expand if only the inferior competitor disperses plastically. This suggests that a trade-off between competitiveness and the ability to plastically adjust dispersal rates can contribute to species coexistence in both static and fluctuating heterogeneous environments. Nevertheless, we also find that coexistence is not possible if the superior competitor has a high maximal dispersal rate but can plastically reduce it at high local resource abundances, thereby evading the conditions for pattern formation and thus the fundamental requirement for coexistence in our case. We propose that this is due to the chosen dispersal cue (resource abundance), which directly translates into fitness (per-capita growth rate) of the consumers. However, dispersal may also be triggered by more indirect cues like crowding (Sutherland et al. 2004; Bowler and Benton 2005; Clobert et al. 2009), which can affect the likelihood of pattern formation and thereby species coexistence.

### 4.2. Relevance of self-organised pattern formation for species coexistence in natural ecosystems

While the structure and parametrisation of our model most closely reflects the properties of plankton organisms in an experimental chemostat setup, we argue that the findings are relevant for many terrestrial and aquatic ecosystems, given the generic structure and general assumptions of our model and the increasing evidence that self-organised pattern formation may be widespread in natural ecosystems (Medvinsky et al. 2002).

Spatial heterogeneity in the biomass distributions of plankton communities is often observed in aquatic ecosystems (Steele 1978) and is predominantly attributed to physical processes such as wind-induced mixing or gradients in light, nutrients and temperature (Platt 1972; Mackas et al. 1985; Abraham 1998). However, a significant share of the observed variation in the biomass distributions of plankton communities cannot be explained by physical processes alone but may rather result from biological processes (Malone and McQueen 1983; Folt and Burns 1999; Kornijów et al. 2020), including self-organised pattern formation (Levin and Segel 1976). A fundamental requirement for this process is that nutrients, autotrophs, and heterotrophs exhibit different diffusion or dispersal rates. In our model, pattern formation occurs for high diffusion rates of nutrients and low dispersal rates of autotrophs. These conditions might be met in very small water bodies or on the microscale within larger water bodies, when molecular diffusion is dominating the movement of nutrients but is irrelevant for macroscopic algal cells (according to the Stokes-Einstein law, Miller 1924). In contrast, in larger, open water bodies the movement of nutrients and phytoplankton is likely to be dominated by transport processes of the water (e.g. convection or turbulent diffusion), implying identical diffusion rates. However, other theoretical approaches have shown that on this scale self-organised pattern formation in planktonic systems may occur based on the different dispersal rates of phyto- and zooplankton (Levin and Segel 1976; Malchow 1993).

A situation that resembles our model system closer are spatially segregated lakes, among which ground water flow may allow for sufficiently fast exchange of nutrients (Hagerthey and Kerfoot 1998; Robinson 2015), while dispersal of phytoplankton is limited to passive transport via wind or animals (Incagnone et al. 2014). In our model, the dispersal rate of the zooplankton not only determines whether pattern formation occurs in the first place, but also which type of patterns emerge. When water bodies are directly connected, active swimming and the more directed movement patterns of larger zooplankton organisms (Dodson and Ramcharan 1991; Pennekamp et al. 2019) may lead to the formation of static biomass patterns, but when zooplankton is restricted to passive dispersal, too, we expect spatio-temporal patterns.

In addition to pattern formation, consumer coexistence in our model further requires that the locally superior competitor has a disadvantage on the regional scale, either by heightened maladaptive switching into resource-poor patches or by insufficient hedging against fluctuating environmental conditions. Evidence exists that especially the second condition is underlying the coexistence of different cladocerans and rotifers. For instance, *Daphnia* species seem to be competitively superior over *Rotifera* species (Gilbert 1985) while suffering more from dispersal limitation between different segregated ponds (Cáceres and Soluk 2002). Similarly, the coexistence of several *Daphnia* species with unstable populations that form metacommunities in small rocky pools appears to result from a trade-off between their competitive abilities and dispersal (and colonisation) rates (Hanski and Ranta 1983).

In terrestrial systems, the autotrophs are sessile plants, i.e. they move only once during their lifetime during seed or propagule dispersal. Their dispersal rate is therefore naturally much smaller than the distribution of nutrients (e.g. via groundwater) or even the active movement of herbivores. While considerable variation in maximal dispersal speeds (Hirt et al. 2017) and strategies (Bowler and Benton 2005) of animals exists, we still expect the comparatively high dispersal rates of terrestrial herbivores to lead to the emergence of static spatial patterns in the species’ biomass distributions. Hence, while vegetation patterns in arid ecosystems are often considered to result from water-mediated scale-dependent feedback (Martinez-Garcia et al. 2022), our results suggest that the feeding pressure by mobile herbivores may also play a role.

### 4.3. Future perspectives

Self-organised pattern formation in our model requires the transport of nutrients, autotrophs, and heterotrophs to occur in both directions along dispersal pathways. This is not always the case in nature as climatic main wind directions or an elevational gradient along freshwater ecosystems can lead to transportation of nutrients and biomass mainly in one direction (Michels et al. 2001; Cottenie et al. 2003). However, theory suggests that spatial pattern formation is still possible under these conditions (Malchow 2000; Brechtel et al. 2018). The preconditions for pattern formation and its effect on species coexistence may further depend on the complexity of natural food-webs, including additional predators and the potential ability of the consumers to plastically increase their emigration rate in response to high predation pressure (Fronhofer et al. 2018).

## Conclusions

We show that two different dispersal strategies known to be able to offset competitive disadvantages, namely avoidance of maladaptive dispersal in static environments and bet-hedging in fluctuating environments, do not require external factors to set the stage for consumer coexistence, but also work if the required spatial heterogeneity emerges due to self-organisation. Compared to externally determined heterogeneity, this process adds dynamic flexibility to metacommunities and creates a novel mechanism for coexistence of consumers based on their different abilities to induce spatial patterns and to benefit from them. At the core of this mechanism, a competitive cycle is generated in which each species affects the environment in a way that allows the other species to recover and to become temporally dominant. Considering that our metacommunity model is very simple and generic, we conclude that spatial instabilities that underlie the demonstrated coexistence mechanisms can be as commonplace as instabilities giving rise to temporal oscillations (e.g. predator-prey oscillations). However, while the latter are often considered as jeopardising species’ persistence, we argue that the former should be interpreted more positively, as they support persistence and enable coexistence.

## Acknowledgements

This study was supported by the German Research Foundation (DFG) under grant number GU 1645/2-1.

## Supporting information

### S1. Flow diagram of the model

In Fig. S1 a flow diagram is provided as a visual representation of the mathematical model, in addition to Eqs. 1 to 7.

**Figure S1:**
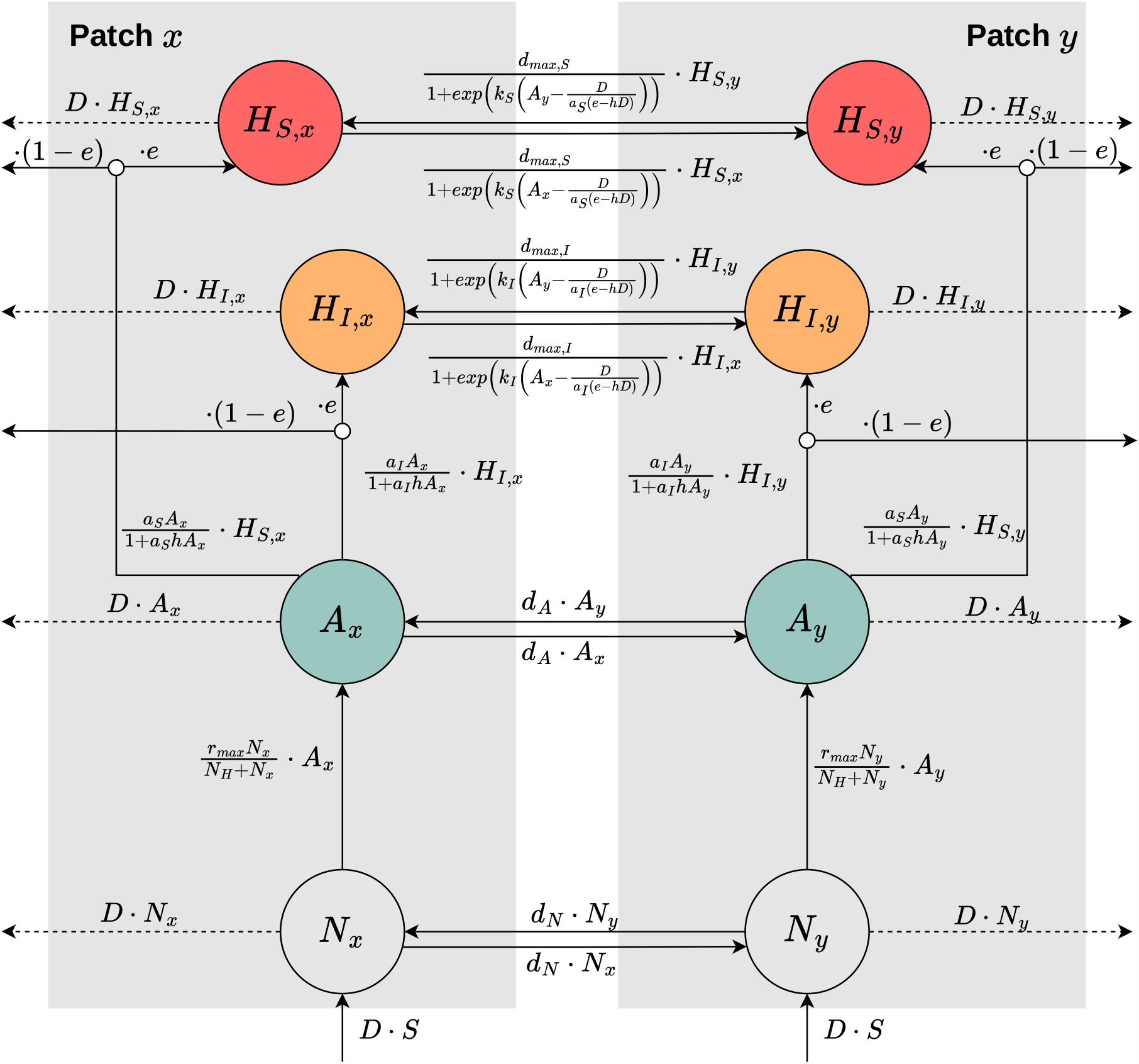
Flow diagram of the different processes contributing to population growth. The state variables (nutrient concentrations *N*, autotroph biomass densities *A*, and biomass densities of the superior and inferior competitor, *H*_*S*_ and *H*_*I*_, all on the two patches *x* and *y*) are in the circles. The total flow rates determining their dynamics are given next to the corresponding arrows connecting the state variables.

### S2. Dynamics in single-patch and spatially homogeneous systems

In spatially homogeneous (i.e., not Turing-unstable) systems with two patches, dispersal does not affect the dynamics, as net dispersal flows between the patches always cancel out. Therefore, the dynamics in this case can be understood by analysing the local population dynamics in a single patch. It is sufficient to do this with just a single heterotrophic consumer present, because the superior competitor always outcompetes the inferior one. Without dispersal, there are no differences between the competitors except for the higher attack rate of the superior competitor, which allows it to feed at a higher rate on the shared resource, irrespective of resource density.

In Fig. S2 a bifurcation diagram of the local system over the attack rate *a*_*H*_ of the heterotroph is shown. The minimal attack rate required for survival of the heterotroph is *a*_*H,inv*_ = 0.48 (invasion threshold). Higher values of the attack rate first lead to stable persistence of the autotroph and heterotroph species, but at *a*_*H,Hop f*_ = 1.33 the system is destabilised in a Hopf bifurcation and predator-prey cycles between the autotroph and the heterotroph emerge. For the results presented in the main text, we deliberately chose attack rates that fall between the invasion threshold and the Hopf bifurcation, but in Appendix S4 we show that the results are not qualitatively affected by the presence of predator-prey cycles in the single-patch (or equivalently, the homogeneous two-patch) system.

**Figure S2:**
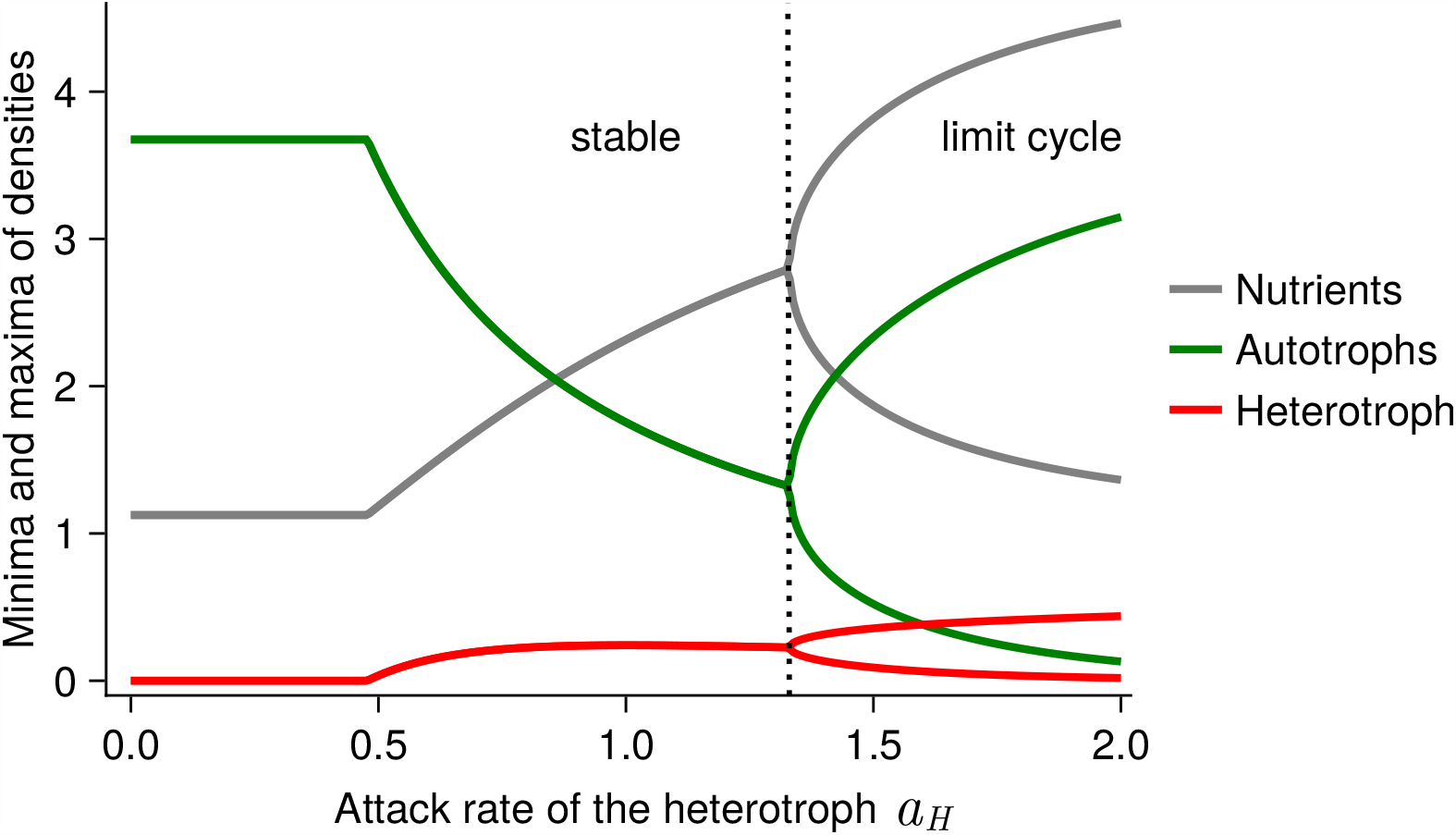
Influence of the attack rate of the heterotroph, *a*_*H*_, on the dynamics of the local system. The Hopf-bifurcation is marked with a vertical dotted line. All parameters as listed in Table 1.

For intermediate values of the dispersal rate of the superior competitor *H*_*S*_, where it does not induce a Turing instability (neither static nor oscillatory), coexistence with the inferior competitor *H*_*I*_ never occurs in the two-patch system, even if *H*_*I*_ induces, when alone, a Turing instability. In Fig. S3 two examples are shown where *H*_*S*_ invades into a system where the resident *H*_*I*_ causes either an oscillatory or a static Turing instability. In both cases, *H*_*S*_ can invade and eventually replace *H*_*I*_, because it has not only the competitive advantage due to its higher attack rate, but also because it has the superior dispersal strategy. When the dispersal rate of *H*_*I*_ is so low that it causes an oscillatory Turing instability (Fig. S3a), the intermediate dispersal rate of *H*_*S*_ allows it to react more flexibly to the fluctuating, heterogeneous resource availability (bet-hedging mechanism). Conversely, when the dispersal rate of *H*_*I*_ is so high that it causes a static Turing instability (Fig. S3b), the intermediate dispersal rate of *H*_*S*_ means less maladaptive dispersal in the emergent source-sink environment. In either case, when *H*_*S*_ becomes more abundant, it dampens the spatial heterogeneity and outcompetes *H*_*I*_ due to its higher attack rate.

**Figure S3:**
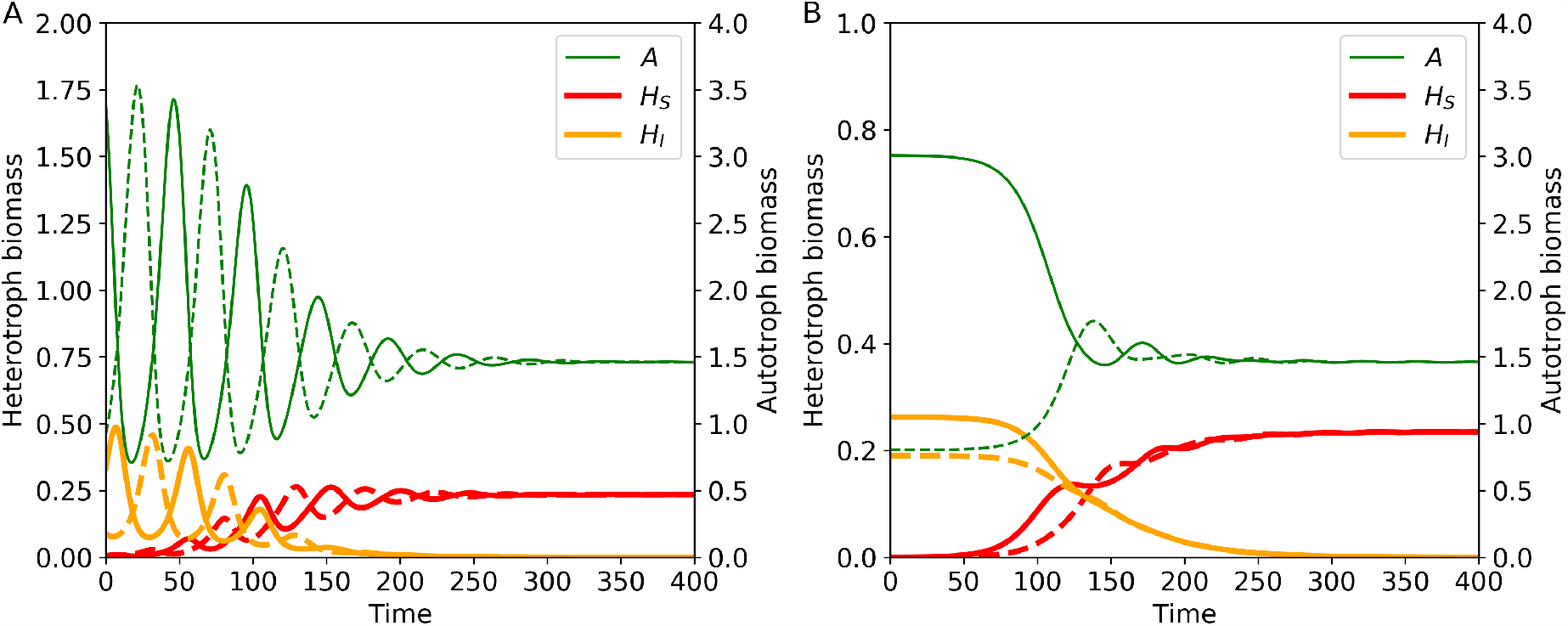
Invasion of the superior competitor *H*_*S*_ into a system where the inferior competitor *H*_*I*_ causes an oscillatory (a) or static (b) Turing instability. Parameters: a) *d*_*max,I*_ = 0.02, b) *d*_*max,I*_ = 0.6, both panels: *d*_*max,S*_ = 0.1, *k*_*S*_ = *k* _*I*_ = 0, all other parameters as in Table 1.

### S3. Criteria for Turing instabilities and ecological mechanisms for pattern formation

Diffusion of a substance in a medium (or likewise, random dispersal of a species in its habitat) usually reduces spatial heterogeneities in its density and over time homogenises the system. However, if multiple substances diffuse (or multiple species disperse) at different rates and interact with each other, a so-called reaction-diffusion system is formed in which positive and negative feedbacks may operate at different spatial scales (scale-dependent feedback, Rietkerk and Van de Koppel 2008). In an ecological context, short-scale facilitation and long-range inhibition, e.g. by spatial redistribution of a limiting resource, may then cause heterogeneous biomass distributions to arise. This process is referred to as self-organised spatial pattern formation because it relies only on the interplay between short-ranged ecological interactions (such as resource consumption and predation) and long-ranged dispersal, rather than on heterogeneous environmental conditions (i.e., any factor that affects the population densities of the species, but is not itself affected by them). In this appendix we first discuss from a mathematical point of view how the conditions for self-organised spatial pattern formation are determined in a system of two discrete habitat patches and then describe in detail the ecological mechanisms leading to the formation of static or spatio-temporal patterns in our model.

#### S3.1. Determining Turing boundaries

The onset of self-organised spatial pattern formation is marked by a bifurcation called Turing instability or diffusion-driven instability. At this bifurcation, a spatially homogeneous equilibrium *X*^*∗*^ = (*N*^*∗*^, *A*^*∗*^, *H*^*∗*)^ becomes unstable. (We here consider a system with only a single heterotroph species *H*, as in the spatially homogeneous system coexistence of two competing heterotrophs is not possible, cf. Appendix S2.) In a system with just two distinct habitat patches as we consider it in this study, the occurrence of a Turing instability can be determined with a simple linear stability analysis (for the general case of network-organised systems with an arbitrary number of patches and dispersal links, see Brechtel et al. 2018). For this, we construct the matrix

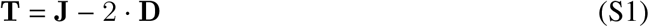

with **J** the Jacobian matrix of the local (single-patch) system evaluated at *X*^*∗*^, **D** the Jacobian matrix of the emigration terms of the system at *X*^*∗*^ (for *k*_*H*_ = 0 this is simply **D** = diag (*d*_*N*_, *d*_*A*_, *d*_*H*_ = *d*_*max,H*_ /2) and the factor 2 as the only non-zero eigenvalue of the Laplacian matrix of the two-patch system. A static Turing instability that leads to the formation of static spatial patterns in nutrient concentrations and population densities of autotrophs and heterotrophs is marked by a single, real eigenvalue of **T** becoming positive, while an oscillatory Turing instability that leads to the formation of spatio-temporal patterns is marked by a pair of complex-conjugate eigenvalues obtaining a positive real part.

If the attractor of the spatially homogeneous system is not an equilibrium but a limit cycle (as is the case if the attack rate *a*_*H*_ *>* 1.33, cf. Appendix S2), one has to resort to Floquet theory (Klausmeier 2008) to determine when this limit cycle is destabilised by the spatial dynamics. In this case, the Jacobian matrix **J** of the non-spatial system has to be evaluated along the limit cycle, which means that the matrix **T** (as above) is now a periodic, time-dependent matrix. Solving the matrix-ODE

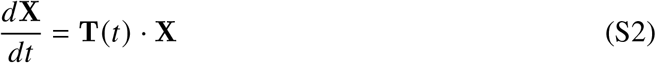

with initial condition **X (**0) = 𝕀 (i.e., the 3 × 3 identity matrix) for exactly one period of the limit cycle, *T*, yields the fundamental matrix **X (***T*). From the eigenvalues of **X (***T*), called Floquet multipliers *ρ*_*i*_, Floquet exponents *μ*_*i*_ = log (*ρ*_*i*_)/*T* can be derived. These Floquet exponents can be interpreted in the same way as the eigenvalues of **T** in the case of a stationary homogeneous equilibrium. Note that this also implies that a static Turing instability of a spatially homogeneous limit cycle leads to the formation of static, not oscillating, patterns in space.

Calculating the zeros of the real part of the dominant eigenvalue(s) of **T** or of the corresponding Floquet multipliers for various combinations of dispersal-related parameters thus reveals the boundaries of the Turing instabilities (Fig. S4). As discussed in the main text, at low values of *d*_*max,H*_, spatio-temporal patterns emerge due to an oscillatory Turing instability, while at high values of *d*_*max,H*_, static patterns emerge due to a static Turing instability. These bifurcations further require that the diffusion constant of the nutrients is above a certain threshold (*d*_*N*_ ≳ 0.1, Fig. S4a) and that the per-capita dispersal rate of the autotrophs is below a certain threshold (*d*_*A*_ ≲ 0.03, Fig. S4b). The further away these parameters are from their respective threshold, the wider are the ranges of values of *d*_*max,H*_ that lead to a Turing instability. Fig. S4c further reveals that the oscillatory Turing instability is not affected by dispersal plasticity, but the static Turing instability is: values of *k*_*H*_ *>* 0.4 prevent this bifurcation completely. Lastly, an example of how the real part of the dominant eigenvalue(s) of **T** changes with *d*_*max,H*_ is shown in Fig. S5 for different values of the attack rate *a*_*H*_. This demonstrates that the parameter ranges under which self-organised pattern formation occurs, increase with increasing *a*_*H*_.

**Figure S4:**
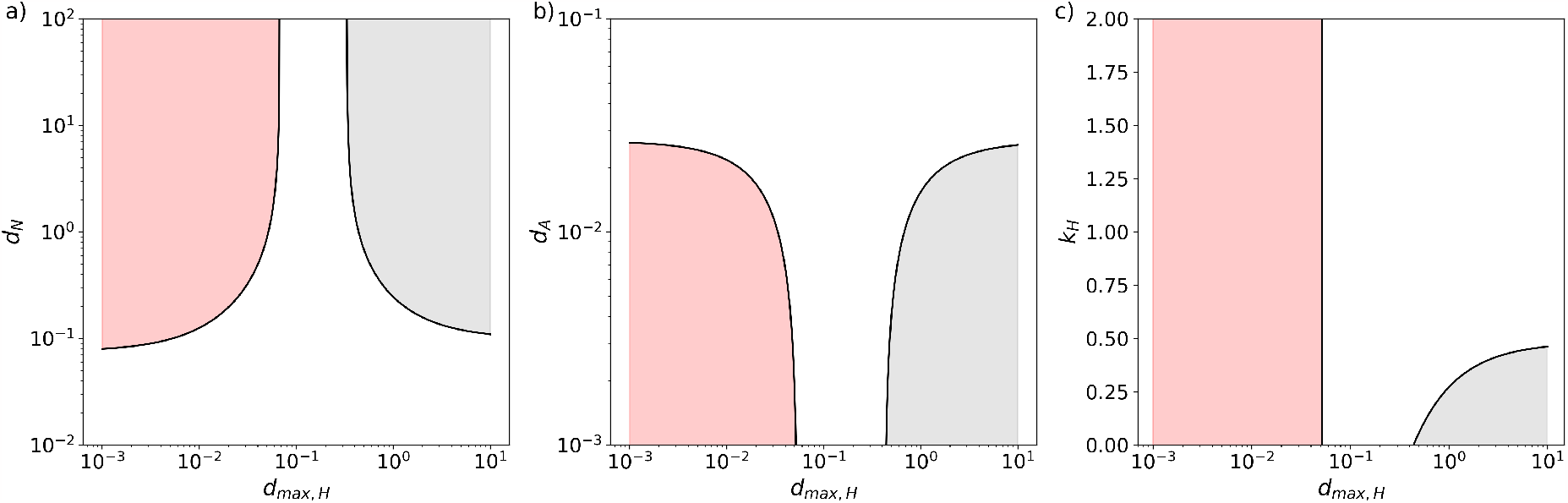
Boundaries of the Turing instabilities in the *d*_*max,H*_ − *d*_*N*_ (a), *d*_*max,H*_ − *d*_*A*_ (b), and *d*_*max,H*_ − *k*_*H*_ (c) planes. In the red-shaded areas spatio-temporal patterns emerge and in the grey-shaded areas static patterns emerge. All parameters (if not varied in the respective panel) as in Table 1.

**Figure S5:**
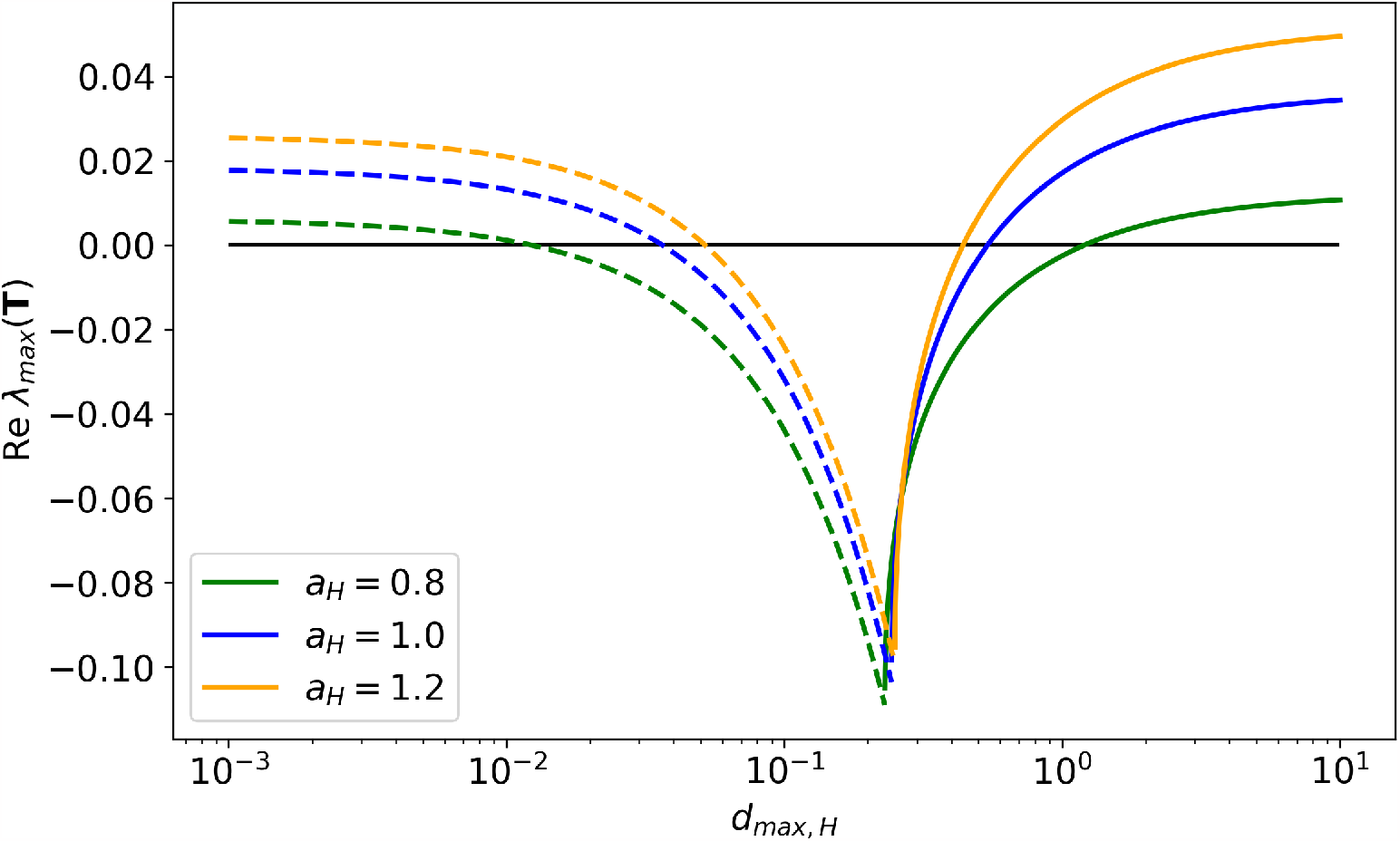
Real part of the dominant eigenvalue(s) of the matrix **T** as a function of the maximal dispersal rate of the heterotroph, *d*_*max,H*_ for different values of the heterotroph’s attack rate *a*_*H*_. Dashed lines: complex eigenvalues, solid lines: real eigenvalues. Parameters: *k*_*H*_ = 0, all other parameters as in Table 1.

#### S3.2. Ecological mechanisms for self-organised spatial pattern formation

##### Destabilisation of the homogeneous equilibrium

Let us first consider a spatially homogeneous equilibrium, i.e., a situation in which nutrient concentrations and biomass densities of autotrophs and heterotrophs are the same on both patches. When the diffusion rate of the nutrients is high, the dispersal rate of the autotrophs is low and the dispersal rate of the heterotroph is either low or high (Fig. S4) this equilibrium is unstable under spatially heterogeneous perturbations.

Any perturbation that favours the autotrophs on one patch (say, patch *x*) relative to those on the other patch (either a direct increase of *A*_*x*_, an increase in nutrient concentration *N*_*x*_, or a decrease in heterotroph density *H*_*x*_) sets the following chain of events in motion: as *A*_*x*_ increases, it suppresses *N*_*x*_ below the level of the homogeneous equilibrium (even if the initial perturbation was an increase in *N*_*x*_). The resulting concentration difference between the patches causes an influx of nutrients from patch *y* which is larger the more *A*_*x*_ suppresses *N*_*x*_. Thus, contrary to expectations from non-spatial systems, a decline of the local nutrient concentration does not stop or even revert the growth of *A*_*x*_, but instead generates additional influx of nutrients that allows *A*_*x*_ to continue to grow (positive short-range feedback). In the other patch (*y*), the outflow of nutrients has the opposite effect as *A*_*y*_ declines (long-range negative feedback). On patch *x*, the increasing autotroph density also allows the heterotroph *H*_*x*_ at least initially to grow, too. The further development, however, depends on whether the heterotrophs have a high or a low dispersal rate.

##### High heterotroph dispersal rate: formation of static patterns

The heterotroph populations *H*_*x*_ and *H*_*y*_ follow the changes in their respective local food availability by increasing or declining, respectively. If their dispersal rate is high, a net flow of heterotroph biomass from patch *x* to patch *y* develops, which prevents the density difference between *H*_*x*_ and *H*_*y*_ from becoming very large. On patch *x, H*_*x*_ can therefore never achieve sufficiently high values to exert effective top-down control of *A*_*x*_ and push it down again, while on patch *y*, the inflated density of *H*_*y*_ puts *A*_*y*_ under severe top-down control. Thus, a high dispersal rate of the heterotrophs cements the system state with low *N*_*x*_, high *A*_*x*_, and moderately high *H*_*x*_ (and vice versa on patch *y*) and a new stable equilibrium with static density differences between the patches emerges.

##### Low heterotroph dispersal rate: formation of spatio-temporal patterns

When the dispersal rate of the heterotrophs is low, no or only a negligible flow of heterotroph biomass occurs as density differences between the patches build up. In consequence, *H*_*x*_ can reach a sufficiently high density to control and eventually suppress *A*_*x*_ below its value at the homogeneous equilibrium.

This causes the nutrient flux between the patches to reverse (now from patch *x* into patch *y*), which allows *A*_*y*_ to grow. The population densities and nutrient concentrations on the two patches thus cycle in anti-phase.

Interestingly, when considering only one patch, the dynamics of autotrophs and heterotrophs resemble classic consumer-resource cycles with *H* following *A* with a delay of a quarter period. However, the cycle period is much longer and the amplitude much larger than what would be observed in a non-spatial system (regular consumer-resource oscillations in a non-spatial version of this model can for example be triggered by increasing the heterotroph’s attack rate *a*_*H*_ beyond 1.33). When spatio-temporal (anti-phase) patterns in the two-patch system emerge, nutrients are constantly shifted back and forth between the patches, which fuels additional growth or decline of the autotrophs, thereby generating the more extreme cycles.

### S4 Sensitivity analyses

#### S4.1. The superior competitor disperses adaptively

Results for a superior competitor with plastic dispersal (*k*_*S*_ = 2) are shown in Fig. S6C-D. The main difference to the case with a randomly dispersing superior competitor (*k*_*S*_ = 0, Fig. S6A-B and Fig. 2 of the main text) is that now no static Turing instability occurs at high *d*_*max,S*_ (cf. Fig. S4C), therefore coexistence is not possible. In contrast, at low *d*_*max,S*_, dispersal plasticity has only a marginal effect on the absolute rate of dispersal.

**Figure S6:**
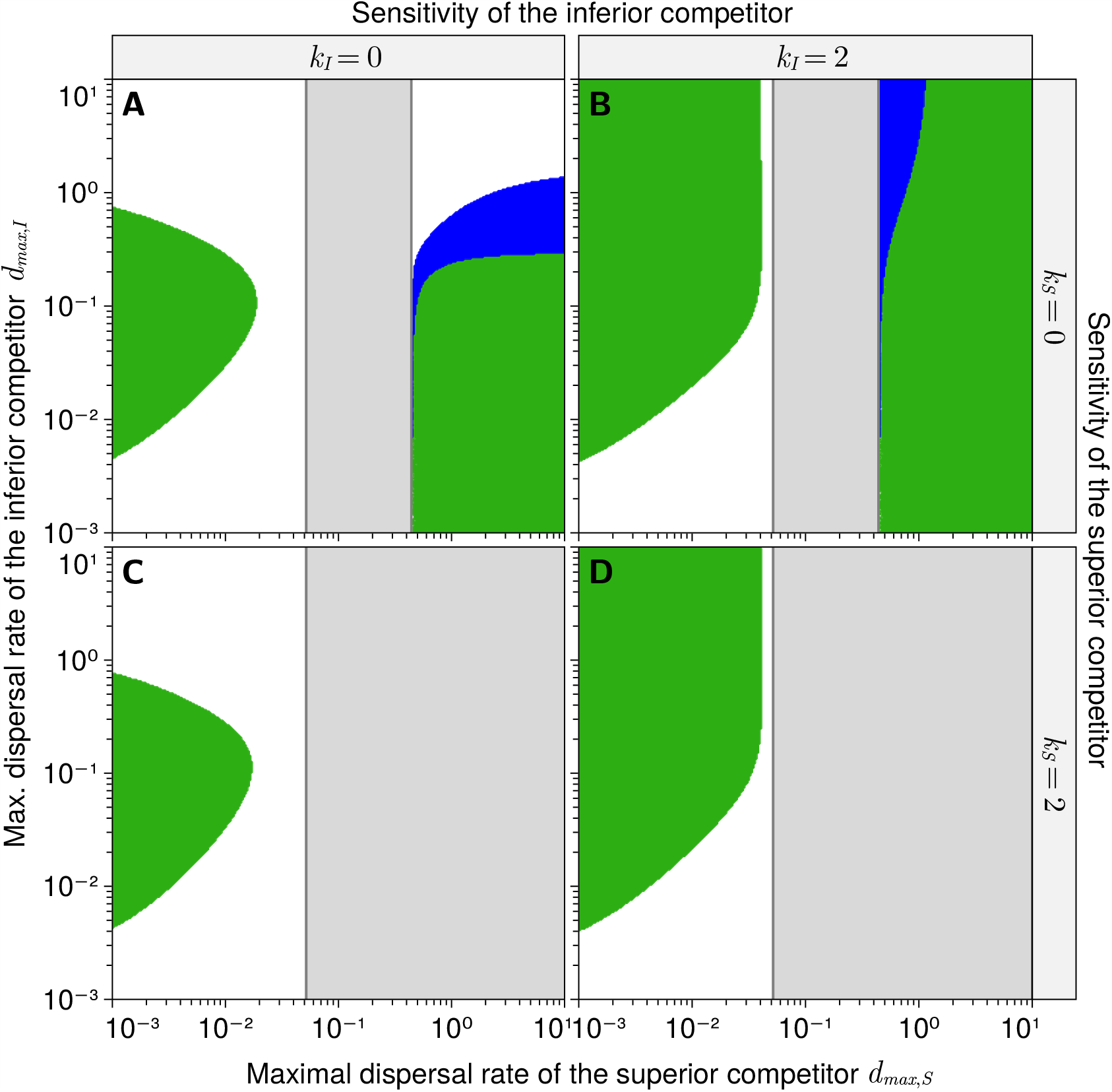
Coexistence of the competitors as function of their maximal dispersal rates. In the green area both competitors coexist with oscillatory dynamics. In the grey shaded area the superior competitor does not induce a Turing instability. The white area denotes the parameter space where a Turing instability occurs, but either the inferior competitor goes extinct. Parameter values, if not provided in the margins of the panels, are as in Table 1.

#### S4.2. Effect of *a*_*I*_ and *k* _*I*_

For three different combinations of maximal dispersal rates of the two competitors we conducted sensitivity analyses to explore the effects of the attack rate of the inferior competitor, *a*_*I*_, and its dispersal sensitivity, *k* _*I*_, in detail (Fig. S7).

**Figure S7:**
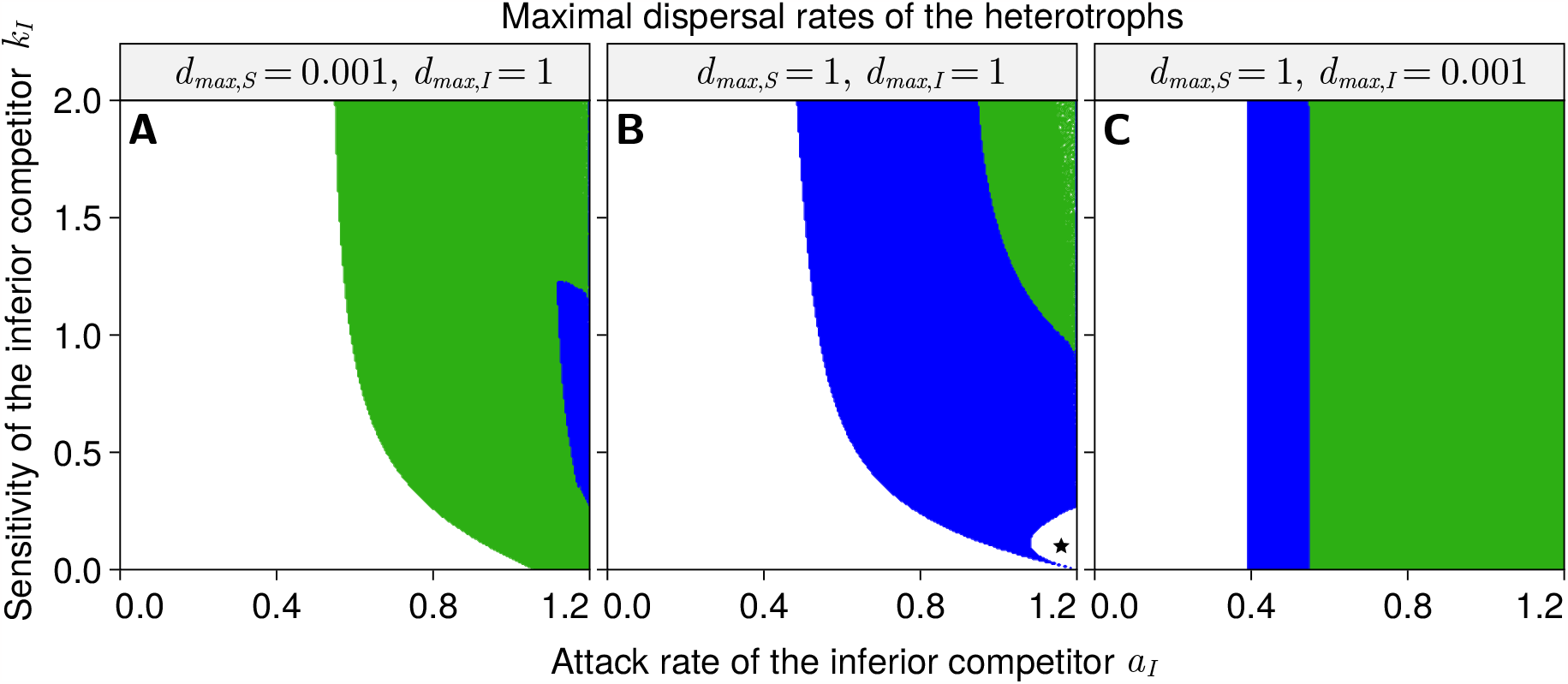
Influence of the attack rate *a*_*I*_ and the sensitivity *k* _*I*_ of the inferior competitor on the dynamics. The superior competitor always disperses randomly (*k*_*S*_ = 0) and the attack rate is set to *a*_*S*_ = 1.2. Both competitors coexist in the regions that are coloured in green (with oscillatory dynamics) and blue (with static dynamics). In the white areas either the inferior or the superior competitor (marked with a star) goes extinct. Parameter values, if not provided in the margins of the panels, are as in Table 1.

In the first case (low dispersal rate of the superior competitor and high dispersal rate of the inferior competitor, Fig. S7A) *H*_*S*_ induces an oscillatory Turing instability and coexistence is made possible by the bet-hedging mechanism. As this is more efficient when *H*_*I*_ can plastically increase its emigration rate from a patch with unfavourable growth conditions and reduce it when growth conditions are favourable, higher values of the sensitivity *k* _*I*_ increase the range of values of *a*_*I*_ for which coexistence is possible.

In the second case (both competitors with a high dispersal rate; Fig. S7B), besides competitive exclusion of *H*_*I*_ or the coexistence of both competitors, we also observe for some parameter combinations the competitive exclusion of *H*_*S*_ by *H*_*I*_. For a large area of the parameter space both competitors coexist without oscillations. If the attack rate *a*_*I*_ and the sensitivity *k* _*I*_ of the inferior competitor increase, coexistence with oscillations occurs. *H*_*I*_ can competitively exclude *H*_*S*_ if *k* _*I*_ is low (but not zero) and *a*_*I*_ is close to *a*_*S*_. This outcome is possible because *H*_*I*_ induces the same type of Turing instability as *H*_*S*_ (in this case a static one, for which *k* _*I*_ must not be too high, cf. Fig. S4C), meaning that it does not lose its advantage due to the superior dispersal strategy once *H*_*S*_ becomes rare.

In the third case (superior competitor with high dispersal rate and inferior competitor with low dispersal rate; Fig. S7C) coexistence of both competitors with or without oscillations occurs. Plasticity of dispersal (i.e., the value of *k* _*I*_) does not play a role at such a low dispersal rate of *H*_*I*_. At low *a*_*I*_, both competitors coexist without oscillations. Here we observe emergent facilitation of *H*_*I*_ by *H*_*S*_: while a single consumer cannot persist if its attack rate is less than 0.48 (Fig. S2), the minimal attack rate for *H*_*I*_ to persist in this case is 0.4. This is possible because the static Turing instability caused by the high dispersal rate of *H*_*S*_ leads to the autotroph biomass being in one patch much higher than its density in the homogeneous system. *H*_*I*_ can take full advantage of this increased food availability in one patch because it only suffers from small emigration losses to the sink patch due to its low dispersal rate. At higher values of *a*_*I*_ the system reveals a very rich dynamic behaviour, as predator-prey oscillations between *H*_*I*_ and the autotrophs on the resource-rich patch emerge. As the amplitude of these oscillations increases, *H*_*I*_ eventually intermittently dominates on one patch and additionally induces an oscillatory Turing instability (at *a*_*I*_ ≈ 0.6). Around this bifurcation point, *H*_*I*_ oscillates with an extreme amplitude, bringing the total population size (*H*_*I,x*_ + *H*_*I,y*_) repeatedly to very low values, thereby creating the risk of stochastic extinction.

#### S4.3. Effect of higher attack rates for both *H*_*S*_ and *H*_*I*_

We further explore the impact of higher attack rates for both competitors on coexistence through pattern formation (Fig. S8). While we kept the absolute difference between *a*_*I*_ and *a*_*S*_ the same, the relative difference between both attack rates is reduced. In addition, the absolute difference between the feeding rates of the competitors at high food concentrations decreases with an overall increase in the attack rates of both competitors. There are two major effects: first, the overall range of dispersal rates that allow for coexistence increases with an increase in the attack rates. Second, we also find three separate parameter regions (two, if *k* _*I*_ = 2) that emerge within the coexistence boundaries where *H*_*I*_ excludes *H*_*S*_. The first, at high dispersal rates for both competitors (*d*_*max,S*_ *> d*_*max,I*_ *>* 0.4), corresponds to the one in Fig. S7B discussed in the previous section, but with a minor difference: while in Fig. S7B *H*_*I*_ avoids maladaptive dispersal through its potential to plastically reduce the dispersal rate, it here does so because it has a constantly lower dispersal rate than *H*_*S*_. In the second region, both competitors have a low dispersal rate (*d*_*max,S*_ *< d*_*max,I*_ *<* 0.04) and exclusion of the superior competitor *H*_*S*_ occurs for the same reason as in the first case: both competitors induce the same type of Turing instability (now an oscillatory one) when they are dominating, meaning again that *H*_*I*_ does not lose its advantage due to the superior dispersal strategy once *H*_*S*_ becomes rare. The third region, at high *d*_*max,S*_ and low *d*_*max,I*_, mirrors the top left corner in panel A. Under the static Turing instability induced by *H*_*S*_, the vastly smaller dispersal rate of *H*_*I*_ is highly beneficial, which allows *H*_*I*_ to dominate and to induce an oscillators instability. A medium-high dispersal rate would allow *H*_*S*_ to come back based on the bet-hedging mechanism, but a very high dispersal rate leads to strong source-sink dynamics over the entire oscillation period, negating the advantage from bet-hedging.

This last region is also interesting because here we observe coexistence under environmental habitat heterogeneity, but competitive exclusion of *H*_*S*_ under self-organised pattern formation (black-and-white pattern in Fig. S8). However, we note that the environmental heterogeneity is based on the (static) biomass patterns induced by *H*_*S*_, which have little in common with the oscillatory patterns observed when *H*_*I*_ prevails. Would the latter be used as basis for the environmental heterogeneity, coexistence would not be possible. A more interesting question is, why coexistence of *H*_*S*_ and *H*_*I*_ under environmental heterogeneity becomes possible in the first place if *d*_*max,I*_ is very small. After all, this should only increase the advantage of *H*_*I*_ in the statically heterogeneous resource landscape. However, as *d*_*max,I*_ gets lower, mortality due to source-sink dynamics decreases. This leads to the emergence of predator-prey oscillations between *H*_*I*_ and *A* on the high resource patch *x* with increasingly large amplitude emerging (Fig. S9). The predator-prey oscillations on the high resource patch also lead to corresponding predator-prey oscillations in the low resource patch, which are in phase with those on the high-resource patch. Eventually, the minimum of *A*_*x*_ during these oscillations drops below the relatively constant (and on average low) value of *A*_*y*_. During these phases, the high dispersal rate *H*_*S*_ allows it to gain a small advantage via the bet-hedging mechanism, enabling it to persist with a small biomass density.

**Figure S8:**
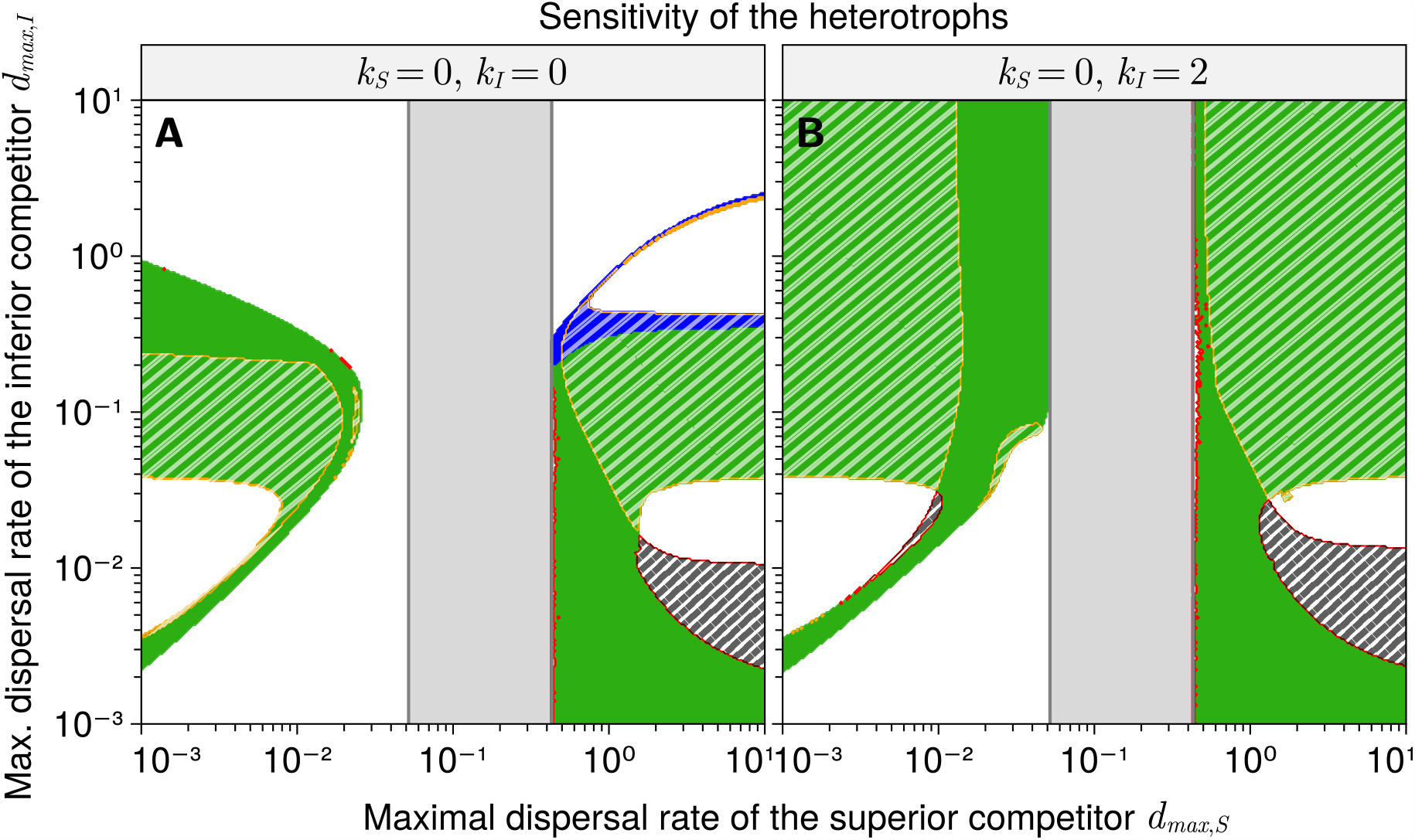
Coexistence as a function of the maximal dispersal rates of the competing heterotrophic species (A: random dispersal of both competitors, B: plastic dispersal of the inferior competitor). Compared to Fig. 2 in the main text, the attack rates of both competitors are larger, but their relative difference is smaller (*a*_*S*_ = 1.5, *a*_*I*_ = 1.3). Self-organised pattern formation allows for coexistence in the coloured regions (blue: static coexistence, green: oscillatory or chaotic dynamics). In the grey area the superior competitor *H*_*S*_ does not induce a Turing instability. In the white-hatched areas the inferior competitor *H*_*I*_ excludes *H*_*S*_ if habitat heterogeneity is determined by the environment instead of emerging in a self-organised way. In the black-hatched areas *H*_*I*_ excludes *H*_*S*_ under self-organised pattern formation, but not if heterogeneity is determined by the environment.

**Figure S9:**
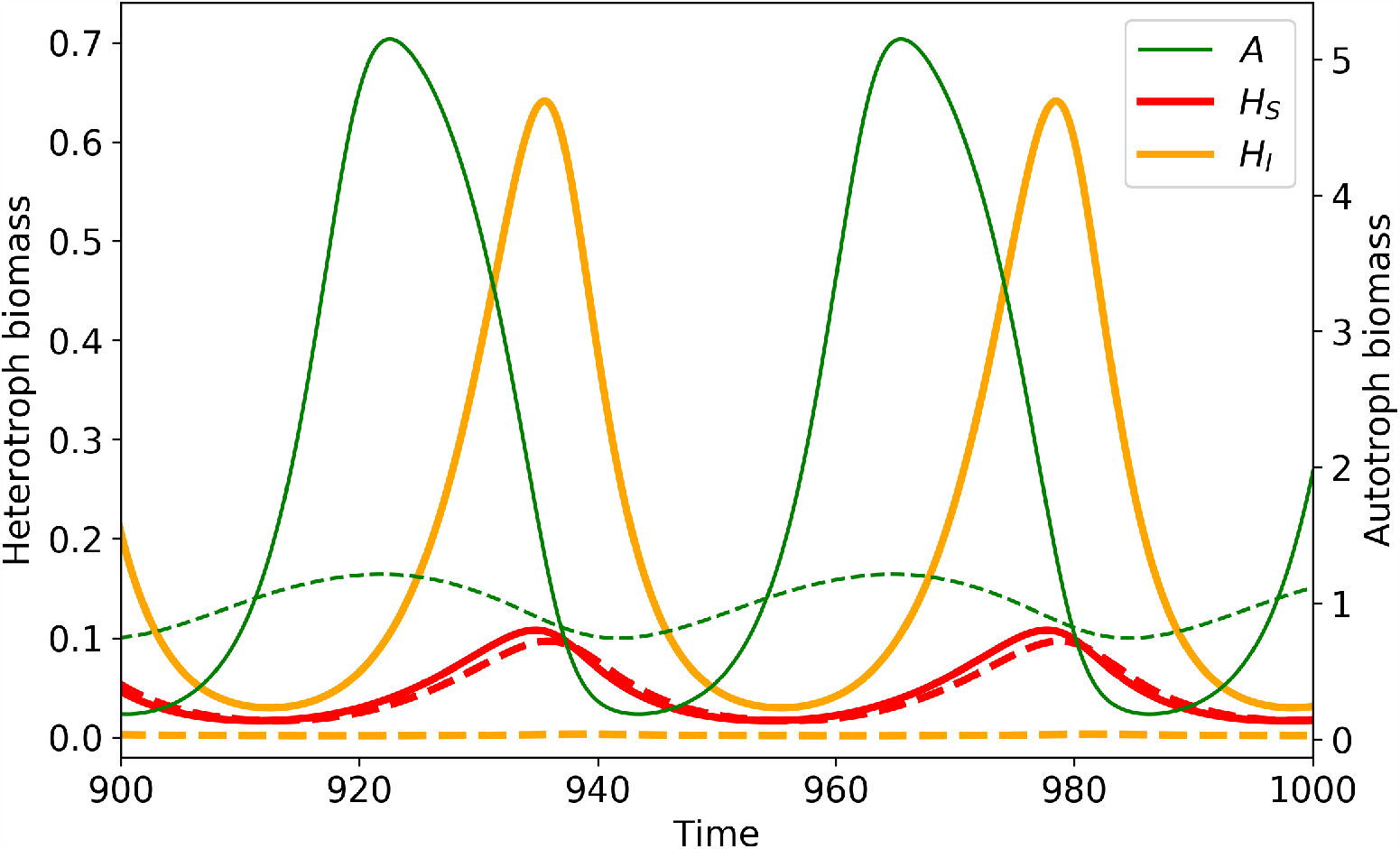
Time series of autotroph (green) and heterotroph biomass densities (*H*_*S*_ in red, *H*_*I*_ in orange) demonstrating coexistence of both competitors under environmental heterogeneity. Solid lines: biomass densities on patch *x*, dashed lines: biomass densities on patch *y*. Parameters: *a*_*S*_ = 1.5, *a*_*I*_ = 1.3, *d*_*max,S*_ = 1, *d*_*max,I*_ = 0.001, *k* _*I*_ = *k*_*S*_ = 0, all other parameters as in Table 1.

#### S4.4. The superior competitor is the invader

In contrast to Fig. 2 in the main text, we here show results for scenarios where the superior competitor *H*_*S*_ is the invader (Fig. S10). Analogously to the procedure for generating the main results, the two patch system was simulated first with only the inferior competitor, *H*_*I*_, for 10^4^ time steps. Then, *H*_*S*_ was added with a low density (*H*_*S,x*_ = 0.001, *H*_*S,y*_ = 0.0001).

**Figure S10:**
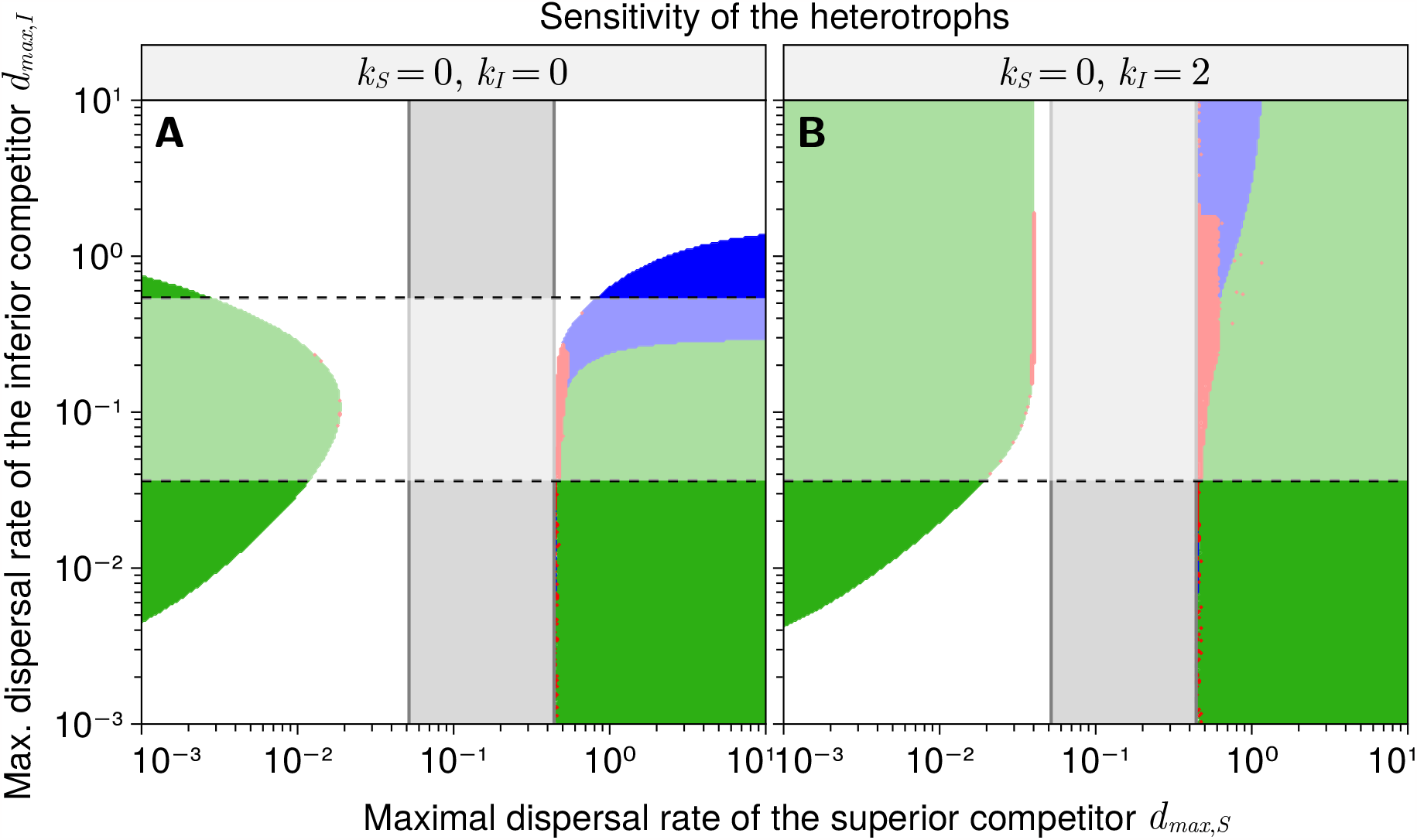
Coexistence as a function of the maximal dispersal rates of the competing heterotrophic species (A: random dispersal of both competitors, B: plastic dispersal of the inferior competitor) in the case of resident *H*_*I*_ and invading *H*_*S*_. Self-organised pattern formation allows for coexistence in the blue and green coloured regions (with static and oscillatory/chaotic dynamics, respectively). Deviations from Fig. 2 are marked in red, here *H*_*I*_ cannot persist. In the grey area the superior competitor *H*_*S*_ does not induce a Turing instability. The white-shaded areas denote where *H*_*I*_ does not induce a Turing instability. All parameters as in Table 1.

In most cases, the results do not depend on the order of invasion of *H*_*S*_ and *H*_*I*_. Only in small regions close to the static Turing instability induced by *H*_*S*_, *H*_*I*_ is excluded if *H*_*S*_ is the invader, but can invade if *H*_*S*_ is the resident (red areas in Fig. S10). In these regions, *H*_*I*_ is not inducing a Turing instability itself (white-shaded areas in Fig. S10), meaning that *H*_*S*_ invades into a spatially homogeneous system where it can easily replace *H*_*I*_ due to its higher competitive strength. Usually, the dominance of *H*_*S*_ leads to the immediate formation of static biomass patterns, which allow *H*_*I*_ to remain in the system. However, this close to the boundary of the Turing instability, the emergence of the patterns takes a very long time, during which *H*_*I*_ continuously declines and eventually has to be considered extinct (Fig. S11A). Since this observation relies on the system balancing for a long time at or near an unstable equilibrium (a situation that would hardly ever occur in nature), we consider it irrelevant for our main conclusions. Also note that once the patterns are fully developed, *H*_*I*_ can of course re-invade the system (Fig. S11B).

**Figure S11:**
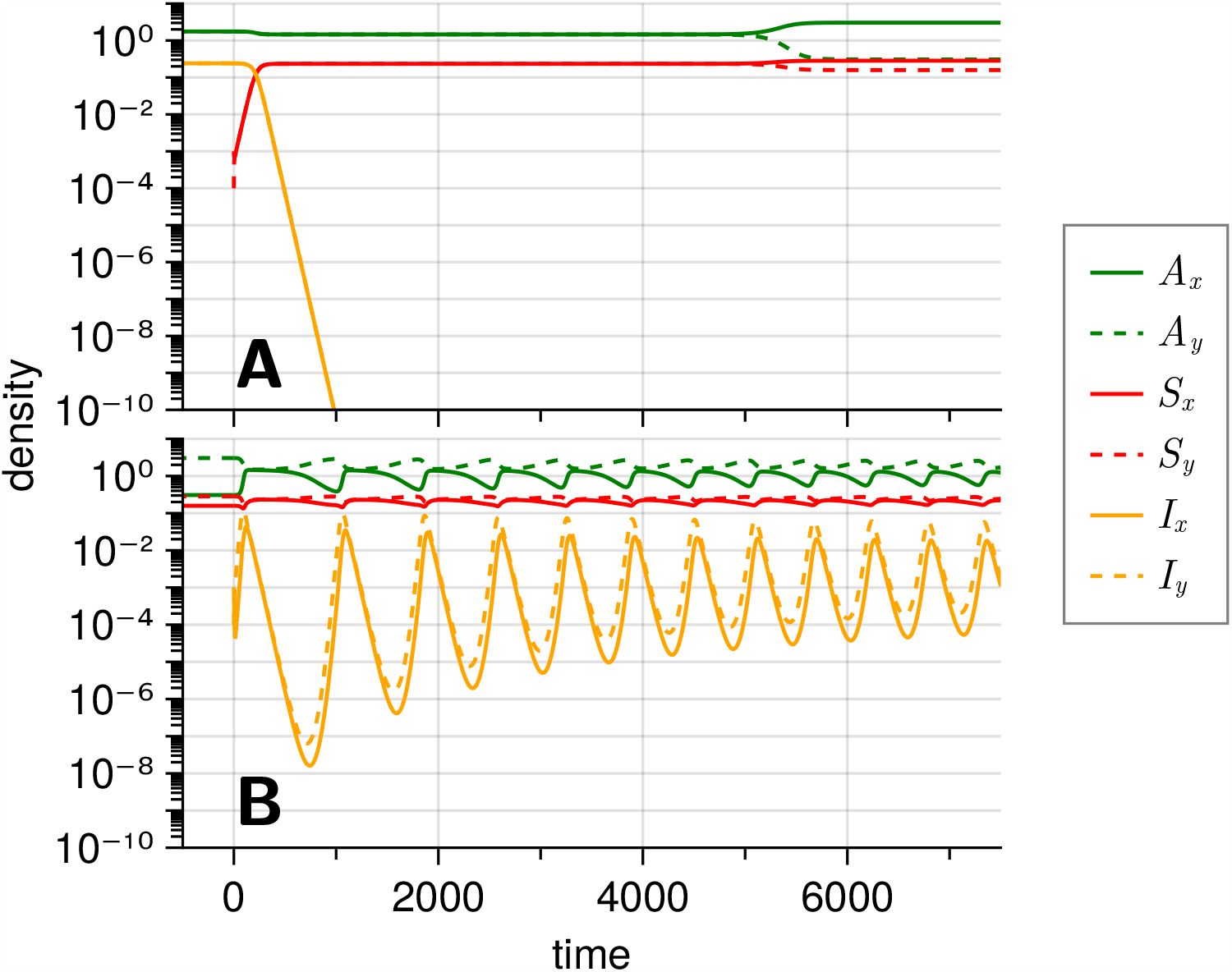
Time series of the autotrophs and the two competitors showing that the competitive outcome can in special cases depend on the order of invasions. In panel A *H*_*I*_ is the resident and *H*_*S*_ the invader, in panel B it is the other way around. Parameters: *k*_*S*_ = 0, *k* _*I*_ = 2, *d*_*max,S*_ = 0.5, *d*_*max,I*_ = 0.2, all other parameters as in Table 1.

### S5. Model with environmental habitat heterogeneity

In this appendix we explain how the model is constructed that includes spatial heterogeneity as environmental factors. Environmental habitat heterogeneity (*eh*) refers to any characteristics of the patches (or rather, difference of characteristics between patches) that affect the dynamics of autotroph and heterotroph populations, but that are not in turn affected by them.

In order to fairly compare the effects of self-organised pattern formation and environmental heterogeneity of the patches we contrast simulations of the standard model, Eq. 1–Eq. 3, (which does not include any environmental heterogeneity between the patches, but allows for self-organised pattern formation) with a model that includes a-priorily as much heterogeneity between the patches as is created by the superior competitor *H*_*S*_ in the standard model, but does not allow for self-organised pattern formation. This *eh*-model is described by the following equations for the dynamics on patch *x*:

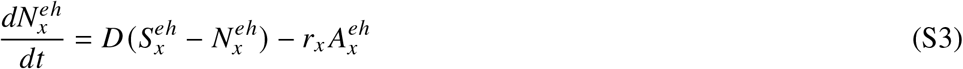

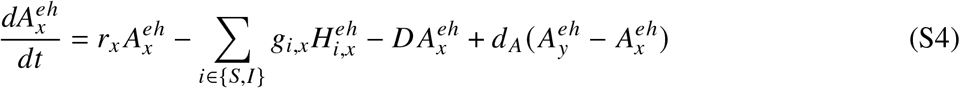

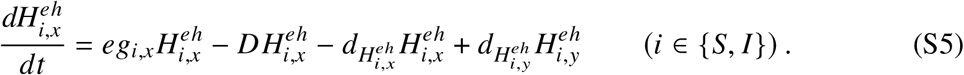

The equations for patch *y* are obtained by swapping the indices *x* and *y*. The nutrient uptake rate of the autotroph, *r*_*x*_, and the grazing rates of the heterotrophs, *g*_*i,x*_ are the same as in the standard model. The key differences between the two models are that in the *eh*-model there is no diffusion of nutrients between the patches (which makes self-organised pattern formation impossible), and that the supply concentrations between the patches differ. We now have

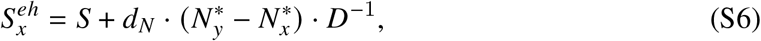

(and equivalently for 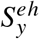), with 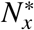 and 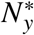 the values of the nutrient concentrations at the attractor of the original model when only the superior competitor *H*_*S*_ is present. Depending on the maximal dispersal rate *d*_*max,S*_ of *H*_*S*_, this attractor is either a limit cycle (at low values of *d*_*max,S*_) with 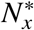 and 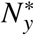 cycling in anti-phase, a homogeneous fixed point at intermediate values of *d*_*max,S*_ (in which case 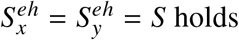) or a heterogeneous fixed point (at high values of *d*_*max,S*_). Through this step, the differences between the patches generated in a self-organised way in the standard model are transferred to the *eh*-model.

When both the standard and the *eh* model are run without the inferior competitor *H*_*I*_, the dynamics they predict are indistinguishable. However, when *H*_*I*_ is present, it can modify the amount of habitat heterogeneity in the standard model, but cannot do so in the *eh*-model. In the main text we have shown that the feedback between emergent habitat heterogeneity and overall competitive abilities of *H*_*S*_ and *H*_*I*_ (including the effect of the respective dispersal strategy) considerably widens the range of conditions under which the two consumer species can coexist. However, this inherent dynamic flexibility also leads to visibly more complex dynamics.

